# Stress testing reveals selective vulnerabilities in protein homeostasis

**DOI:** 10.1101/2025.06.11.659168

**Authors:** Berent Aldikacti, Hidayet Putun, Vishal Sarsani, Rilee Zeinert, Patrick Flaherty, Peter Chien

## Abstract

Protein quality control (PQC) systems are essential for cellular resilience to proteotoxic stress. Despite intensive study for decades, functional redundancies in the system obscure the contributions of the collectively important individual genes. Here, we leverage transposon sequencing across bacteria strains lacking key chaperones and proteases to reveal hidden determinants of stress response in protein homeostasis. By profiling fitness under multiple proteotoxic stresses, we uncover stress-specific vulnerabilities and reveal how major players of PQC mask correlations between transcriptomic responses and gene fitness. As an illustration of unexpected connections, we identify a heat-specific synthetic lethality between the disaggregase ClpB and DNA Polymerase I (PolA) mediated by persistent aggregation of the RecA recombinase and toxic persistence of the heat shock regulon. Our findings reveal that stress-induced aggregation is not broadly toxic. Rather, it becomes lethal in specific genetic or environmental contexts due to the depletion of components only needed in those specific circumstances. This work presents a framework to reveal normally hidden fragility in stress responses using gene fitness scores adaptable to a variety of systems.

## Introduction

Protein quality control (PQC) is critical for cell homeostasis across all life. Networks of molecular chaperones and proteases balance protein synthesis, folding, maturation, and degradation to maintain a healthy proteome. Disruption of human PQC results in several pathologies, such as neurodegenerative diseases^1,2^, cancer^3^, and aging^4,5^. In bacteria, loss of PQC components affects normal cell growth, stress responses, and host interactions^2^.

Mutagenesis screens have been widely used to identify the phenotypic consequences of genetic perturbations. Methods such as transposon insertion sequencing (Tn-seq) or CRISPRi have been used to identify the phenotypic profiles of single-gene knockouts or knockdowns on a genome-wide scale. These approaches have led to the discovery of new virulence factors^6,7^, membrane integrity elements, motility genes^8^, and whole-genome essentiality maps of several bacteria^9,10^. However, due to the crosstalk between pathways and redundancies in biological systems, most genes do not show phenotypic consequences in single-gene knockouts^11,12^.

Simply put, while a gene may be necessary for a stress response, deletion of only this gene may not result in an observable phenotype because redundancy in the networks will compensate for this loss. Consequently, numerous genes remain either “hypothetical” or annotated based on structural similarity, even in organisms that have been extensively studied and have well-annotated genomes^13^. To reveal the functions of genes whose function is obscured by redundancy, there have been recent developments, such as dual CRISPRi-seq^14^ and dual Tn-seq^15^, to interrogate gene-gene interactions on a genome-wide scale. Additionally, many gene phenotypes can only be revealed during stress. For example, the disaggregase ClpB does not show any phenotype under normal growth conditions; however, it is essential under acute heat stress^16–18^.

We reasoned that combining known PQC stresses under conditions where known response factors are compromised would reveal insight into protein homeostasis akin to failure mode analysis in engineering, which reveals the robustness of a system once specific components have been weakened. Specifically, we interrogate genome-wide fitness of Tn-seq libraries generated in strains missing major chaperones and proteases under known proteotoxic stress conditions **(Figure 1A)**. Through this approach, we identify genes that are important to the global stress response, robust against perturbations in quality control pathways, and sensitive to perturbations in specific PQC pathways. We identify redundancies in PQC that normally mask the correlation of transcriptional and phenotypic consequences of stress response genes. Finally, we identify an unexpected synergy between heat stress-induced protein aggregation and the DNA damage response, which exposes how cells become vulnerable during heat stress due to specific environmental or genetic pressures rather than a generic defect in fitness. Taken together, our work serves as an exemplar of combining systems-wide fitness surveys with a deliberate focus on specific stresses to reveal novel insights.

**Figure 1.**
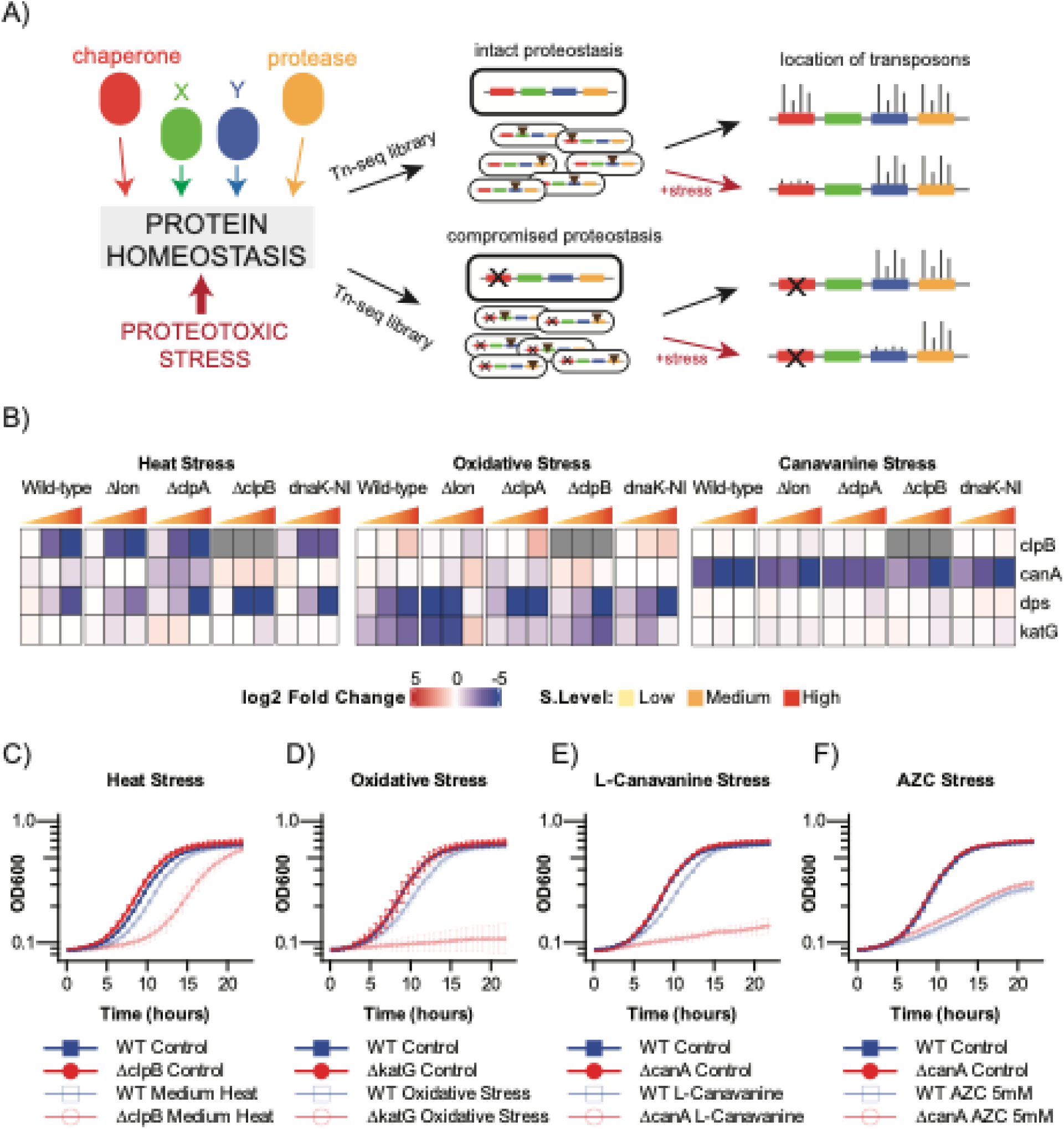
Global proteotoxic stress fitness determinants in *Caulobacter crescentus*. **(A)** Maintaining cellular proteostasis requires the contribution of chaperones, proteases and other proteins. Transposon insertion libraries result in single disruptions in every cell (middle), the position of which can be deconvoluted through sequencing and number of insertions quantified (right). In wildtype cells, the importance of major genes during specific proteotoxic stress responses is revealed (top: shows that red chaperone gene is important for stress tolerance, while blue protein X is not). Generating these libraries in cells compromised in proteostasis due to loss of a factor (red chaperone in this case) and subjected to the same proteotoxic stress unmasks the hidden players in the response (blue protein X is important in the absence of red chaperone protein in this illustration) **(B)** Heatmaps showing fitness scores (log2 fold change) of genes from −5 (blue) to +5 (red) range for the mutant library independent stress determinants. Fold changes are calculated by subtracting the log2 batch-corrected transposon insertion count of the stress condition from that mutant library control condition. ClpB insertions under Δ*clpB* strain greyed out to indicate there are no insertions in *clpB* gene in that strain. **(C, D, E, F)** Bacterial growth curves (OD600) showing mean +- standard deviation from three biological replicates of (C) *WT* and *ΔclpB* strains under medium heat stress (42°C 45 minutes), (D) *WT* and Δ*katG* strains under medium oxidative stress (0.05mM H_2_O_2_, chronic), (E) *WT* and Δ*canA* under medium l-canavanine stress (50 µg/ml, chronic), and (F) *WT* and Δ*canA* under azetidine-2-carboxylic acid (AZC) stress. (5mM, chronic). All experiment performed as three biological replicates and OD600 measured every 20 minutes. Dots and error bars plotted every 40 minutes for visualization purposes.

## Results

### Phenotypic fitness profiling for studying protein quality control network

Protein quality control (PQC) stress responses rely on the presence of chaperones and proteases that mitigate proteotoxicity. Major players of this stress response are known, such as Hsp70 chaperones and disaggregases, that are the primary responders to the misfolded protein burden^19^. How additional players shape this response is less known in part because of the dominance of these major players. We reasoned that a multiplexed reverse genetic screen approach such as Tn-seq conducted in cells lacking major PQC responders and subjected to proteotoxic stresses would identify fitness determinants that may be hidden or redundant. While PQC systems range in components across systems, the core of the PQC system is preserved in bacteria. Given this conservation we used *Caulobacter crescentus* (*Caulobacter*) in our current approach.

We previously generated dense transposon mutagenesis libraries from five strains, including loss of proteases (Lon, ClpA) and loss of normal chaperone activity (ClpA, ClpB, DnaK)^20^. We subjected these mutagenesis libraries to three proteotoxic stressors at three different stress levels: heat stress, which causes misfolding, degradation, and DNA damage; oxidative stress, which oxidizes proteins and causes misfolding and DNA damage^21,22^; and L-canavanine, an L-arginine analog that leads to protein misfolding upon incorporation into polypeptides. A Bayesian analysis of this dataset exposed the overall structure of the data, showcasing synergistic connections between strains and stress conditions, but lacked specific molecular insights due to the nature of this initial global analysis^20^.

To explore this dataset in a more candidate-focused manner, we first identified genes that affected fitness for a given stress in all strains **(Figure 1B).** For example, ClpB is the Hsp104-family ATP-dependent disaggregase needed to resolve heat-induced protein aggregates^18,23^. As expected, *clpB* is identified as a fitness determinant in all strains, specifically under heat stress, which we validated by constructing clpB deletions in all strains and performed growth assays of these double mutants (**Figure 1C, Supp. Fig. 1**). Similarly, KatG is the only known peroxide-detoxifying catalase in *Caulobacter. W*e find that *katG* is identified as a fitness determinant in all strains, specifically under oxidative stress **(Figure 1D, Supp. Fig. 2)**.

For canavanine stress, we find *CCNA_02154* (renamed CanA) as a fitness determinant in all strains, which we validate using double mutant strain growth curves (**Figure 1E, Supp. Fig. 3**). We considered if increased sensitivity to canavanine was due to excess incorporation of this specific unnatural amino acid or toxic consequences from prolific levels of aberrantly folded polypeptides. In agreement with the first possibility, loss of *canA* does not increase sensitivity to azetidine-2-carboxylic acid (AZC), a proline analog that is also misincorporated and leads to proteome-wide misfolding (**Figure 1F**). Given that CanA is predicted to be an acetyltransferase and its selective effects on canavanine, we hypothesized that acetylation of canavanine may prevent its incorporation into polypeptides. Consistent with this model, CCNA_02153, the gene adjacent to *canA*, is annotated as a deacetylase, and based on Tn-seq, loss of this gene protects against canavanine toxicity (**Supp. Fig. 4**).

The genes *clpB, katG,* and *canA* are all examples of determinants that are universally important for one stress response (heat, oxidative, and canavanine, respectively) but do not show differences in fitness for the other tested proteotoxic stresses. We found additional genes that show this same degree of universal stress specificity (**Supp. Table 1**). We also identify examples of genes being important for more than one response (**Supp. Table 2**). For example, *dps* (*CCNA_02966*) encodes for the ferritin-like DNA binding protein and emerges as a fitness determinant for both oxidative and heat stress in our studies, consistent with work from other bacteria^24,25^. We validated the dual stress intolerances of *dps* in *Caulobacter* by constructing double mutants and find sensitivity to both peroxide and heat in all strains (**Supp. Fig. 5**).

Finally, we tested if any genes were fitness determinants in all stresses and all strains. While no genes passed this filter, we identified 18 fitness determinants for all proteotoxic stresses tested in several strain combinations (**Supp. Table 3**). This indicates that no global fitness determinants are important for all tested PQC stresses that function entirely independently of the chaperones and proteases examined in this study.

### Loss of Lon highlights synergies between DNA damage and oxidative stress responses

Given the depth of the dataset, we next explored subsets of genes known to be regulated by proteotoxic stresses, focusing on the known transcriptional response to oxidative stress^26^. In wild-type cells, we found that most genes that are transcriptionally upregulated during oxidative stress do not contribute to tolerance of that stress **(Figure 2A)**. A similar, originally surprising, lack of correlation between transcriptomic and fitness profiles of genes has been reported in other systems^12,27^, leading to a natural conclusion that genes needed for stress tolerance should already be expressed at levels needed to withstand that stress prior to the insult. Interestingly, we find that a subset of these upregulated genes, which includes DNA damage repair proteins, are revealed as important for oxidative stress tolerance in the absence of the Lon protease **(Figure 2B**). Given that oxidative stress is also genotoxic, an interpretation of this result is that DNA damage repair is critical once cells have dealt with the initial insult of the stress. This result reflects the strength of our approach using multiple strains, as this link was only evident in cells lacking the Lon protease **(Figure 2C**).

**Figure 2.**
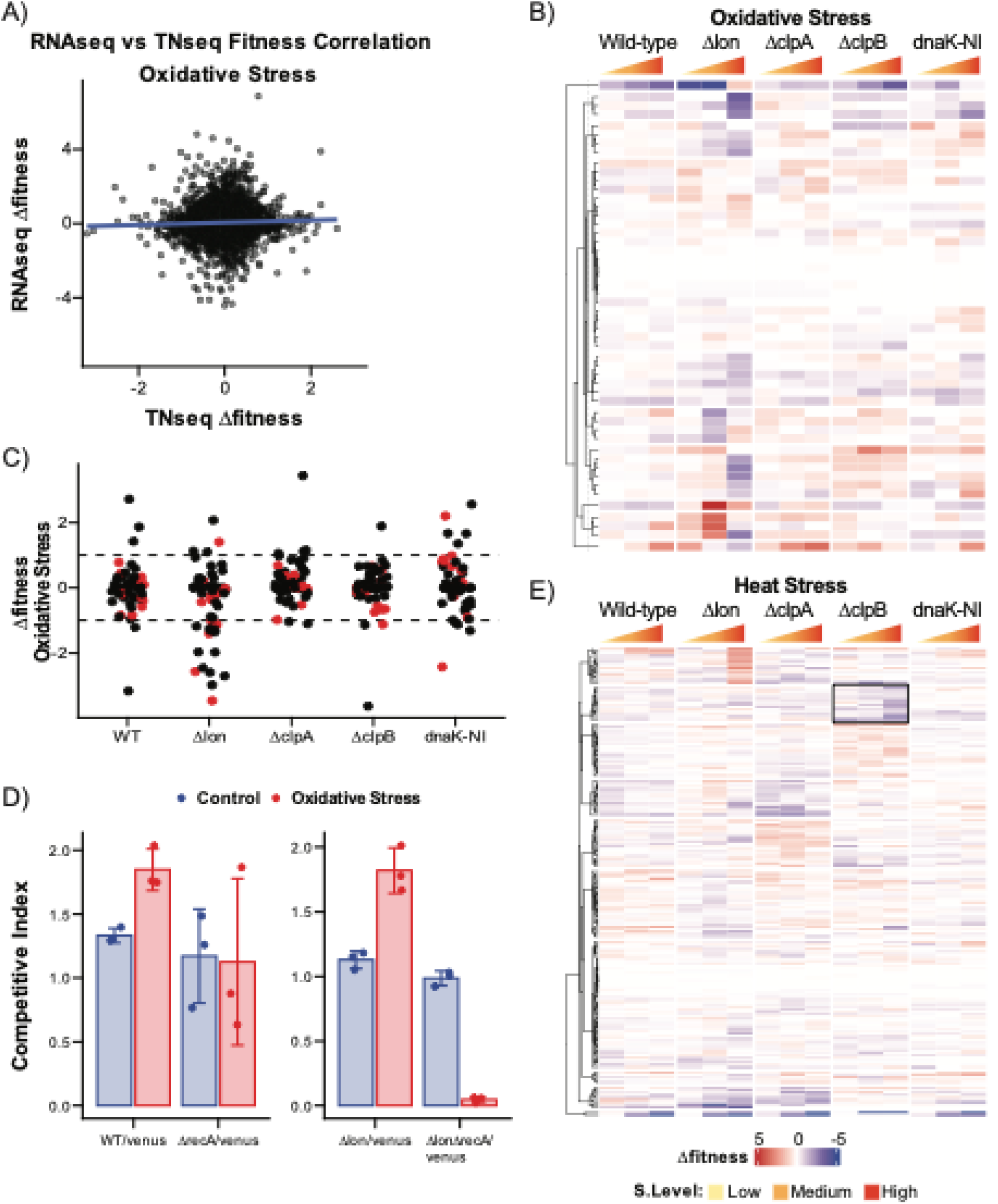
Loss of Lon protease uncovers synergism between oxidative stress and DNA damage. **(A)** Scatter plot compares the log2 fold change from *WT* oxidative stress transcriptomic data and *WT* oxidative stress Tnseq data. Blue line indicates the linear fitted line to all of the data. Transcriptomic data is obtained from Silva et al. The *WT* 15-minute H_2_O_2_ condition compared to the *WT* control. **(B, C)** Oxidative stress regulon obtained from Silva et al^26^. and used to compare transcriptomic changes to fitness. The differentially expressed genes in the *WT* 15-minute H_2_O_2_ condition compared to the *WT* control. (B) Heatmap shows the log2 fold changes of batch-corrected transposon insertion counts (Δfitness) for low, medium, and high oxidative stress conditions for *WT, Δlon, ΔclpA, ΔclpB*, and *dnaK-NI* transposon libraries. (C) Fitness values from high oxidative stress are plotted as dots to highlight the distribution between strains. Dashed lines indicate fitness at −1 and +1. Red dots highlight the DNA damage response genes identified by Modell et al^67^. **(D)** In vivo competition experiments for *WT, ΔrecA, Δlon*, and *ΔlonΔrecA* strains under 0.025 mM H_2_O_2_ stress were conducted, with the competitive index as the output parameter. The experiment involved mixing the strains *WT-WTvenus*, *ΔrecA-WTvenus*, *Δlon-Δlonvenus*, and *ΔlonΔrecA-Δlonvenus* in a 1:1 ratio and growing them for 24 hours. Image quantification of the ratio of fluorescent to non-fluorescent cells was performed at the time of mixing and after 24 hours of growth to calculate the competitive index. Each dot represents a biological replicate, and the error bars indicate standard deviation. **(E)** Heatmap shows the log2 fold changes of batch-corrected transposon insertion counts for low, medium, and high oxidative stress conditions for *WT, Δlon, ΔclpA, ΔclpB*, and *dnaK-NI* transposon libraries. RpoH regulon obtained from Schramm et al^28^. and used to compare transcriptomic changes to fitness. The black box highlights the cluster of genes that cause a negative fitness change in the *ΔclpB* background. This cluster includes the genes *yaaA* and *oxyR*.

We validate the link between Lon, oxidative stress, and the need for DNA repair proteins by performing competition growth assays. RecA is the recombinase critical for many aspects of DNA damage and is one member of the Lon-specific oxidative stress-sensitive gene sets (**Figure 2B,C**). We generated cells lacking *recA* or *lon* or both, then fluorescently tagged strains with mVenus to visualize populations of different strains upon coculture growth. We find that loss of RecA does not affect competitive growth compared to wildtype cells following oxidative stress, but Δ*lon* strains lacking RecA do worse than Δ*lon* cells alone under stress conditions **(Figure 2D, Supp. Fig. 6)**. In addition, monitoring the growth of monocultures of each strain, *ΔlonΔrecA* show a synergistic sensitivity to oxidative damage. We conclude that upregulation of DNA damage response genes is an important aspect of the oxidative stress response, but a Lon-dependent pathway normally masks its phenotypic importance.

Similar analysis with other transcriptional responses, such as the heat stress regulon controlled by RpoH^28,29^, also exposes the importance of specific genes in conditional backgrounds with greater numbers of genes emerging as important during heat stress in specific strains **(Figure 2E, Supp. Fig. 7)**. For example, the oxidative stress genes *CCNA_03496* (*yaaA)*^30,31^ and *CCNA_03811* (*oxyR)*^26^ emerge as important for heat stress tolerance, specifically in a *ΔclpB* background, but not other strains, suggesting a link between heat and oxidative stress when ClpB is lost that differs from other strains **(Supp. Fig. 8-9)**. Our prior work supports that Δ*clpB* strains are altered in oxidative stress response during low heat stress compared to wild-type strains^20^, consistent with the selective gene fitness in this current analysis. Similar analyses will provide a rich starting ground for experiments to test new hypotheses regarding the PQC system in bacteria.

### Loss of DNA polymerase 1 and ClpB are synthetically lethal during heat-stress

During our examination of strain-specific stress responses, we noticed that DNA polymerase 1 (*polA; CCNA_03577*) resulted in both oxidative and heat stress sensitivity in our datasets. However, in a ΔclpB background, the heat stress sensitivity was markedly stronger than the oxidative stress sensitivity **(Figure 3A),** and we decided to follow up on this observation. PolA is the first discovered DNA polymerase^32^ and plays a crucial role in DNA replication and repair, mainly due to its function in the formation of Okazaki fragments^33^. In *E. coli*, upregulation of *polA* is part of the DNA damage (or SOS) response, and deletion of *polA* results in sensitivity to DNA damage, heat, and oxidative stress^34,35^. We identify similar defects for a Δ*polA* strain of *Caulobacter* (**Figure 3B-D**). Consistent with the Tn-seq results, loss of *polA* in a Δ*clpB* background resulted in a marked synergistic sensitivity to heat stress **(Figure 3C, Supp. Fig. 10).**

**Figure 3.**
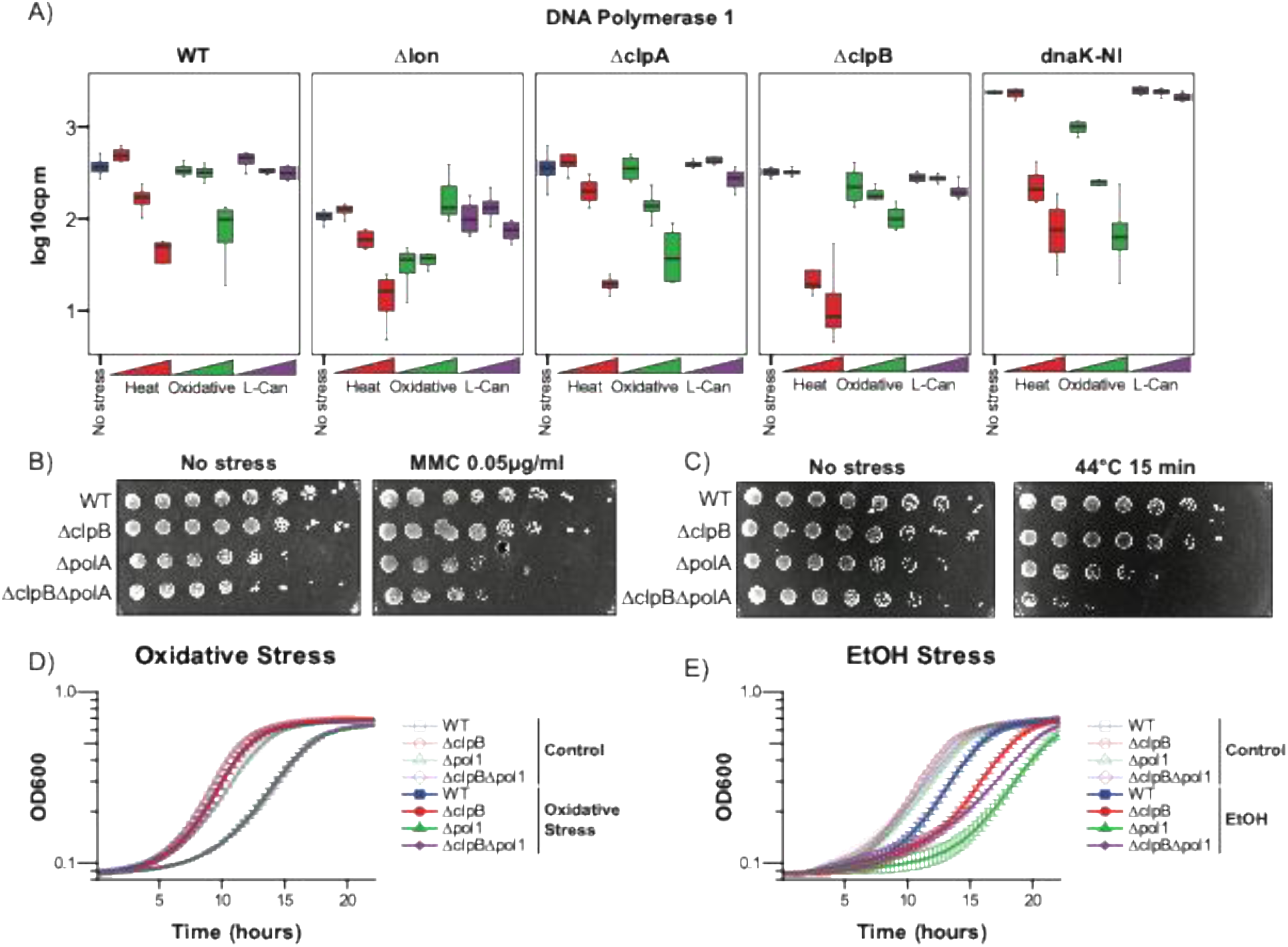
DNA polymerase 1 is sensitive to heat and oxidative stresses and shows synergistic fitness defect with ClpB under heat-stress. **(A)** DNA polymerase 1 (PolA) transposon insertion profile across all tested transposon mutagenesis libraries and stresses. Insertion counts are presented as log10 counts per million after batch correction. Stress conditions are color-coded, depicting low, medium, and high stress levels through the triangle. Bars show the median and standard deviation of the insertion counts. **(B)** Spot titration (10-fold dilution series) of *WT, ΔclpB, ΔpolA*, and *ΔclpBΔpolA* strains under 0.05 µg/ml chronic Mitomycin C (MMC) stress. **(C)** Spot titration (10-fold dilution series) of *WT, ΔclpB, ΔpolA*, and *ΔclpBΔpolA* strains under 44°C heat stress for 15 minutes. **(D)** Growth curves (OD600) *of WT, ΔclpB, ΔpolA*, and *ΔclpBΔpolA* strains under chronic 0.05mM H_2_O_2_ stress. **(E)** Growth curves (OD600) of *WT, ΔclpB, ΔpolA*, and *ΔclpBΔpolA* strains under %20 EtOH stress for 15 minutes.

Given that PolA is known to be important for other stresses such as DNA damage and oxidative damage, we also tested genotoxic and oxidative sensitivities in the double mutants and found no synergy with loss of ClpB (**Figure 3B&D**). Similarly, other defects of Δ*clpB*, such as sensitivity to ethanol^18^, were not enhanced in the absence of PolA (**Figure 3E**). We conclude that the synergistic fitness defect we find between ClpB and PolA is exclusive to heat stress and sought to understand if this was due to a systems-level or molecular-level defect.

### Persistent upregulation of the heat-stress regulon is toxic in the absence of PolA

To explore a systems-level basis of this synergistic fitness defect, we used RNA sequencing to compare the transcriptomic profile of wildtype and Δ*clpB* strains during moderate heat stress conditions. We found 72 genes up-regulated, and 45 down-regulated in Δ*clpB* under moderate heat stress (FDR < 0.05; **Supp. data 1)**. We reasoned that synthetic fitness effects could arise from lower expression of factors needed in a Δ*polA* background or intoxication due to overexpressed factors. Of the downregulated genes, we focused on the *imuABC* operon, which encodes a mutagenic polymerase, because this operon, along with other SOS response genes, is also upregulated in the Δ*polA* strain based on RNA-seq (**Supp. Fig. 11, Supp. data 1**), suggesting its loss may explain the synthetic sickness of the double mutant. However, deletion of *imuC* (also known as *dnaE2*^36,37^) in a Δ*polA* background shows no synthetic phenotype even with heat stress (**Supp. Fig. 12**). We, therefore, turned our attention to the genes upregulated during heat stress in Δ*clpB*.

In bacteria, transcriptional changes due to heat stress are primarily driven by the sigma factor RpoH^38–40^. Prior studies in *Caulobacter* suggest that ClpB is needed for reactivation of the housekeeping sigma factor RpoD following heat stress^18^, which would result in the eviction of the unstable heat shock sigma factor RpoH from the RNAP complex during the recovery phase. Consistent with this interpretation, we found that nearly all of the up-regulated genes in Δ*clpB* strains can be explained by the RpoH-dependent regulon^28^ (**Figure 4A**). We considered whether persistent expression of the heat stress regulon may underlie some of the synthetic lethality we see between ClpB and PolA during heat stress. Upregulation of RpoH is well tolerated in wild-type cells, but expression of a stabilized active variant of RpoH (RpoH-V56A) results in lethality. Interestingly, in cells lacking PolA, even the upregulation of wild-type RpoH causes lethality (**Figure 4B**), suggesting that Δ*polA* strains are particularly sensitive to excess RpoH activity.

**Figure 4.**
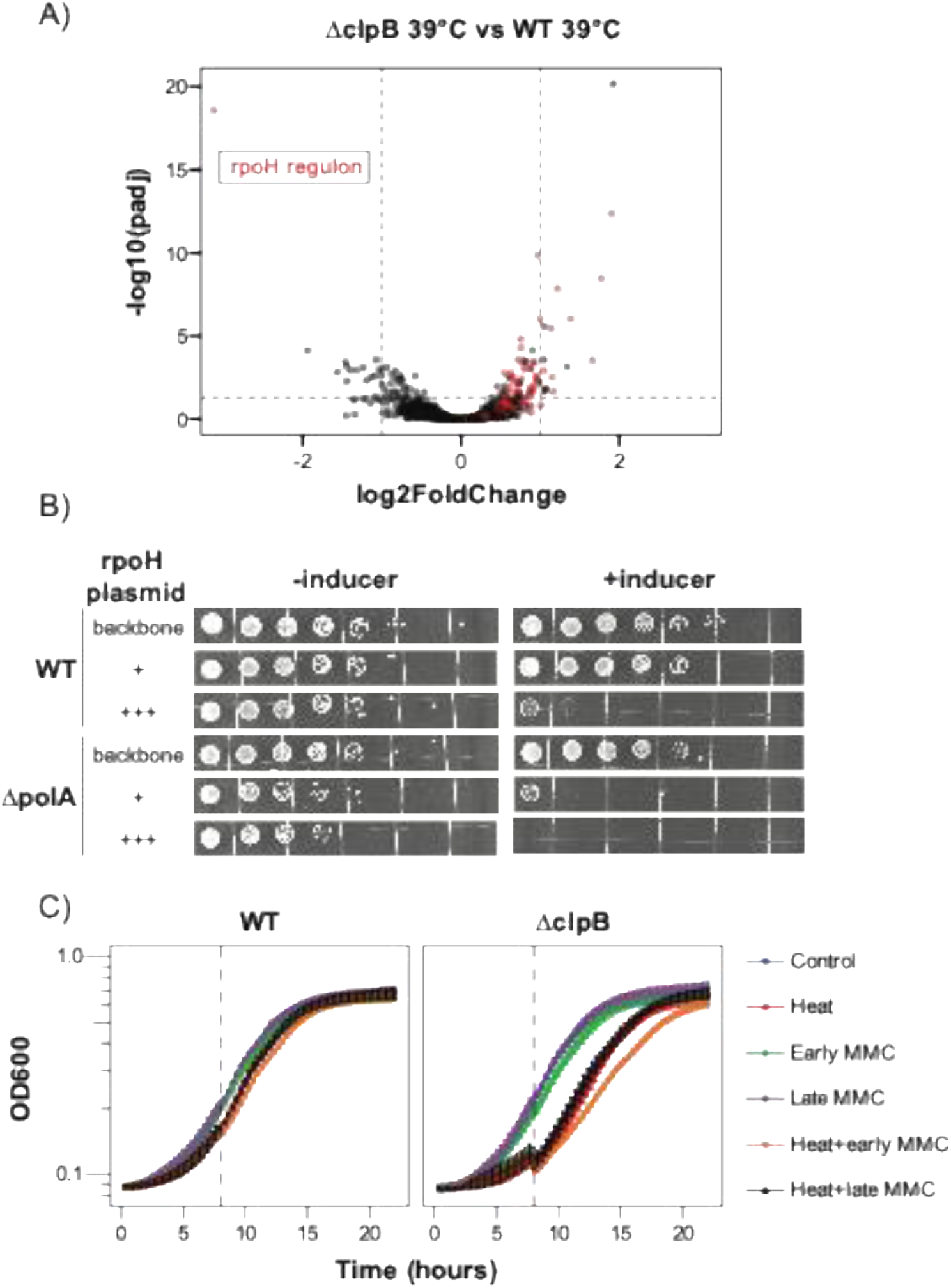
Consistent *rpoH* expression contributes to synergistic PolA ClpB heat stress phenotype. **(A)** Total RNA sequencing was performed to identify transcriptomic differences between Δ*clpB* and *WT* strains under heat stress. (39°C, chronic) rpoH regulon^28^ highlighted on the volcano plot in red. **(B)** Spot titration (10-fold serial dilutions) with *WT* and Δ*polA* strains with rpoH overexpression plasmids. Vanillate (0.5mM) used as the inducer, and chloramphenicol (1µg/ml) used as the selection. The experiment ran as biological replicates, and representative image shown. **(C)** Growthcurves (OD600) of *WT*, and Δ*clpB* strains under Heat (44°C 15min), Early MMC (added at 0h, 0.05 µg/m, chronic), Late MMC (added at 8h, 0.05µg/ml, chronic), and combinations of Heat and early or late MMC conditions. Dashed line indicates the late MMC addition at 8 hours. Experiments done in biological triplicates, and results shown in mean +- standard deviation.

Because the SOS regulon is persistently upregulated in Δ*polA* and the heat-shock regulon is persistently upregulated in Δ*clpB*, we wondered whether transcriptional conflicts between these regulons might contribute to the PolA-ClpB phenotype. To do test this, we primed Δ*clpB* cells with UV stress to induce the SOS response and then exposed them to heat stress. In contrast to a model where elevated SOS responses conflict with the heat stress regulon, our results indicated that activation of the SOS response provided protection against heat stress **(Supp. Fig. 13).** This suggests that persistent upregulation of the SOS response in Δ*polA* cells does not contribute to the synergy between PolA and ClpB that results in reduced heat stress tolerance.

### Loss of ClpB results in sensitivity to immediate DNA damage upon heat stress

Given the known genotoxic sensitivity of Δ*polA* and the synergy between PolA/ClpB, we explored links between DNA damage and ClpB-mediated protection from heat stress. Interestingly, cells were more sensitive to the DNA-damaging agent Mitomycin C (MMC) immediately after heat stress but lost this sensitivity following an 8-hour recovery after heat stress. This effect was more pronounced in the Δ*clpB* strain (**Figure 4C**), suggesting that the failure to recover after heat stress and increased sensitivity to immediate DNA damage might be linked. Because RpoH-dependent expression is persistent in Δ*clpB* (**Figure 4A**) during moderate heat stress, we tested whether this persistence could be the root cause of the DNA damage sensitivity. We found that cells expressing RpoH variants were no more sensitive to DNA damage than cells with control plasmids (**Supp Fig. 14**). Therefore, while elevated RpoH-dependent expression explains some features of the Δ*clpB* defects, it is insufficient to explain this immediate DNA damage sensitivity. Because ClpB is primarily known as a disaggregase, we explored whether aggregation of specific proteins could underlie this phenomenon.

### Heat-induced aggregation of specific proteins contributes to the PolA-ClpB phenotype

Prior work identified several proteins aggregated during heat stress in *Caulobacter*, including GyrA and RecA^41^, both important for DNA replication and repair. In *E. coli,* genetic studies suggest that *polA* and *recA* are synthetically lethal^42,43^. Building on this, we hypothesized that the sequestration of RecA in insoluble protein aggregates that might contribute to the synergistic effect between PolA and ClpB, as cells lacking ClpB will fail to resolve their aggregates. This hypothesis would require that *ΔpolA* and Δ*recA* show synthetic fitness effects. Initial attempts to generate double mutants failed, which is consistent with a need for both genes. We were able to construct Δ*polA*Δ*recA* strains in the presence of an integrated copy of *recA* at a separate locus under inducible control. Removal of the inducer results in cell death, which is consistent with a synthetic lethal effect due to the loss of both RecA and PolA (**Figure 5A**). Armed with this knowledge, we next explored if this synthetic lethality underlies the synergistic defect of Δ*polA*Δ*clpB* during heat stress.

**Figure 5.**
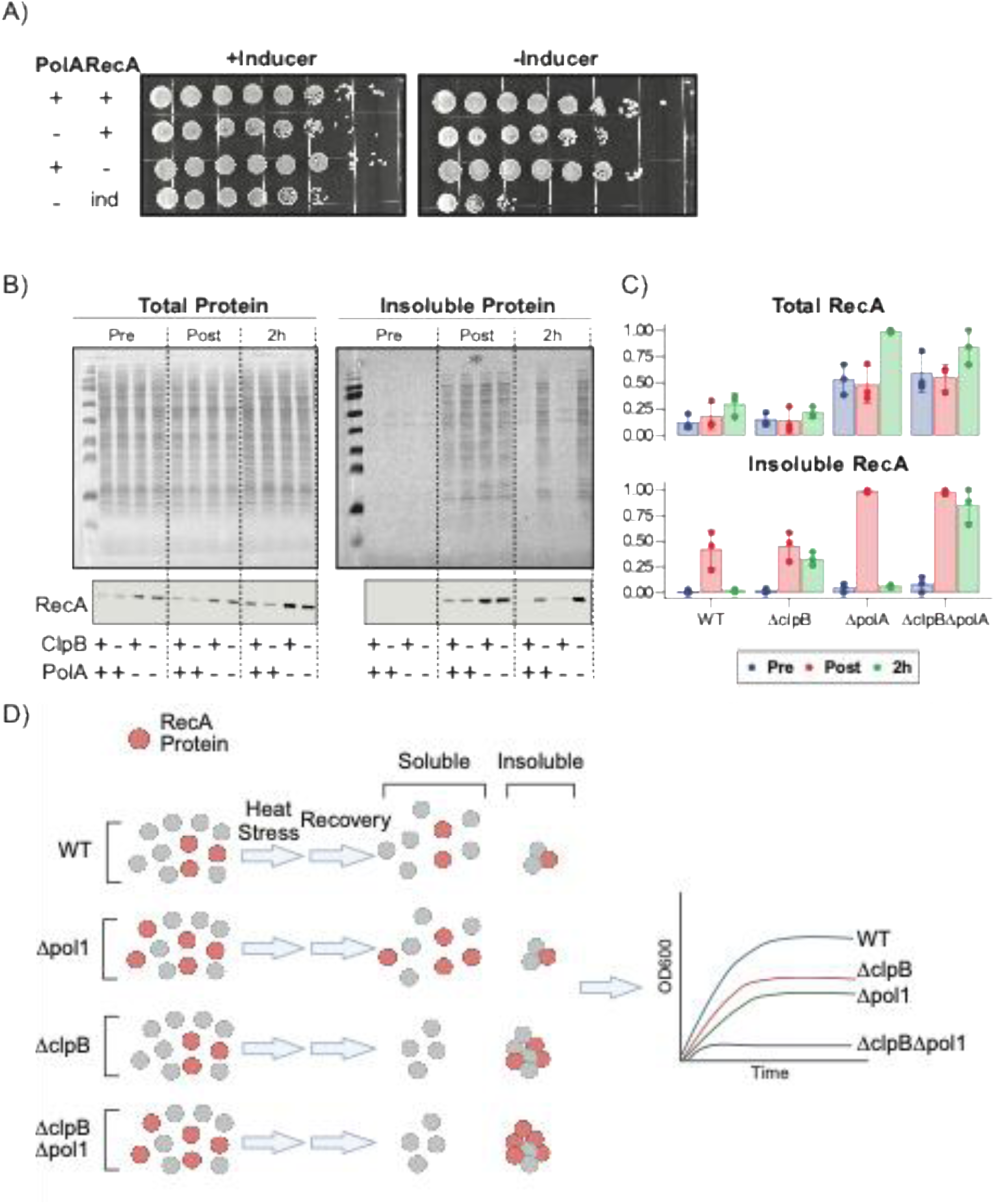
Heat-induced aggregation of RecA in Δ*clpB* causes increased sensitivity to heat stress in the absence of PolA. **(A)** Spot titrations (10-fold serial dilutions) with *WT* or cells lacking RecA or PolA or both in an inducible RecA background. “+/−” indicate the presence or absence of PolA or RecA, and “++” strain indicates the *WT* strains. %0.2 Xylose was used as the inducer, and %0.2 Glucose was used to suppress the leaky expression of the Xylose promoter in the “-Inducer” condition. “ind” in the legend indicates inducable RecA. **(B)** Total and insoluble protein fractions collected from *WT, ΔclpB, ΔpolA*, and *ΔclpBΔpolA* strains before (pre), after (post), and 2 hours (2h) after heat stress. Ponceau stain is used to identify total protein (upper panel) and western blots for RecA proteins. (Lower panel) Quantifications were performed by normalizing RecA levels to the total protein and shown in the right panel as bar plots. Representative blots are shown. Experiments performed in biological triplicates. **(C)** RecA western blot quantifications are shown from Figure 5A (lower panel). Dots represent individual biological replicates, and error bars represent standard deviation. **(D)** Cartoon schematic of how RecA aggregation in ΔclpB strains leads to depletion of RecA from the soluble pool.

We measured insoluble RecA levels during heat stress using centrifugation of extracts before heat stress and during recovery, then detected RecA with specific antibodies. In Δ*polA* strains, soluble RecA levels were elevated under basal conditions, consistent with an increased need for this protein in the absence of DNA polymerase I and the increased transcription of the SOS response (**Supp. Fig. 11**). As predicted, acute heat stress causes aggregation of many proteins, including RecA (**Figure 5B-C**). In cells with wildtype ClpB activity, these aggregates were resolubilized over the time course, while in the absence of ClpB, aggregates were persistent. Western blots confirmed that RecA was also retained in aggregates in the absence of ClpB (**Figure 5C)**. We conclude that prolonged loss of soluble RecA in the Δ*polA*Δ*clpB* strain following heat stress contributes to the heat stress specific synthetic lethality in this double mutant.

Given that Δ*clpB* strains show persistent aggregation of many proteins but are more fit than Δ*polA*Δ*clpB* strains following heat stress, we propose that heat stress-induced aggregation alone is not itself toxic. Instead, the aggregation of particular proteins (such as RecA) is toxic only under certain genetic backgrounds (such as Δ*polA*) or additional stresses (such as MMC) that make the strain under those conditions vulnerable to that specific loss. Given this rationale, other heat-stress-driven synthetic lethalities only in Δ*clpB* might reflect these additional vulnerabilities. For example, the importance of oxidative stress factors YaaA and OxyR for tolerating high heat stress in Δ*clpB* described earlier (**Supp. Fig. 8-9**) may be due to the aggregation of redundant proteins important for the oxidative stress response that fail to be rescued in the absence of ClpB.

## Discussion

This study demonstrates how targeted transposon sequencing can reveal critical insights into protein quality control (PQC) mechanisms in proteotoxic stress conditions. By perturbing PQC pathways and subjecting cells to various stressors, we identified new genetic determinants of stress resilience and uncovered condition-specific vulnerabilities. Specifically, we identify new genes important for stress responses, demonstrate how redundancies in the stress response mask the importance of important genes, and demonstrate that loss of viability due to aggregation results from condition-specific vulnerability rather than general aggregation (**Figure 6**).

**Figure 6.**
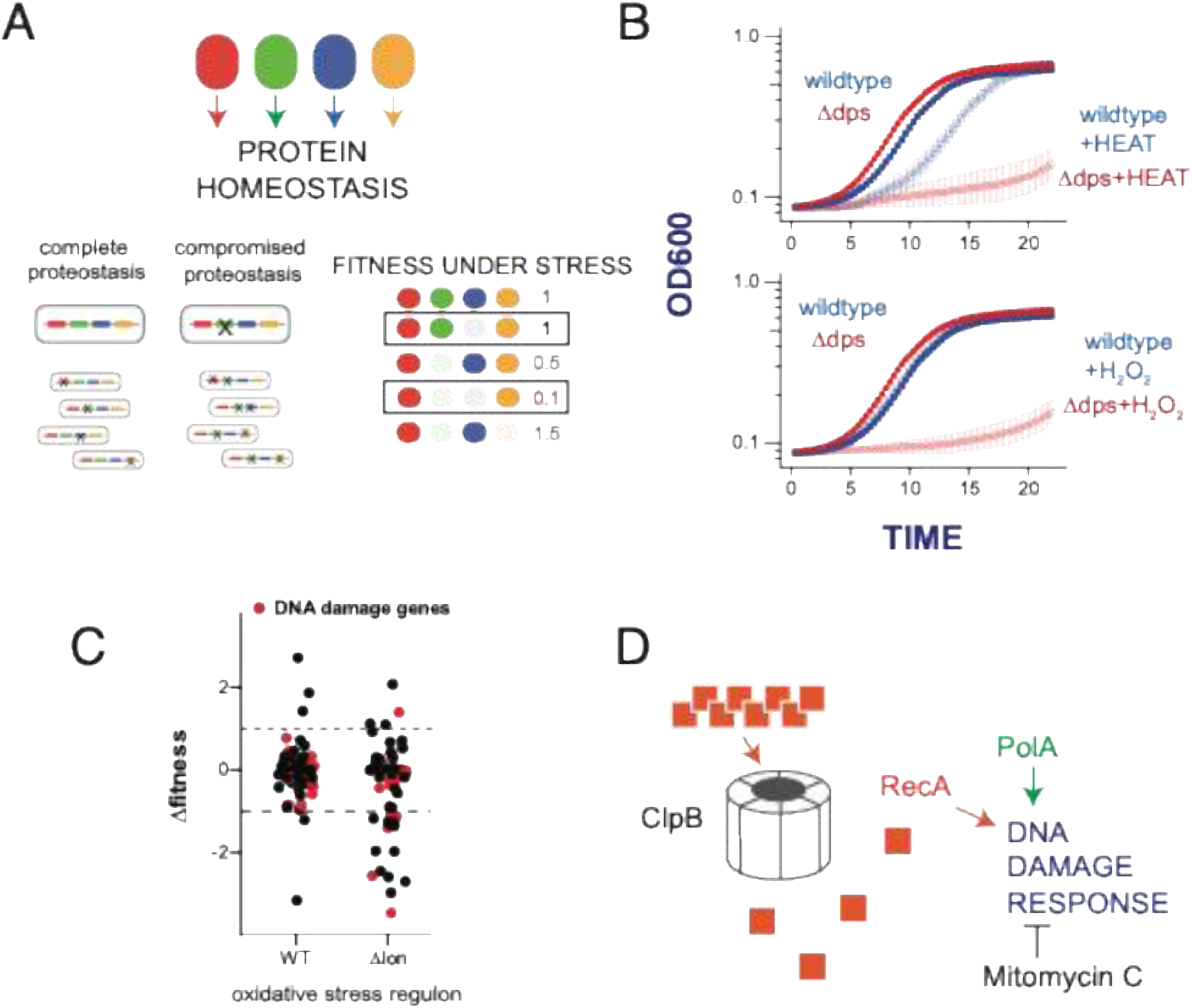
**(A)** Unmasking vulnerable genes arise from use of strains deficient in PQC components under environmental conditions that specifically stress the protein homeostasis. This approach unearths new genes that affect stress responses **(B)**, correlations in phenotypes and transcription under vulnerable conditions **(C)**, and deepens understanding of intersections of heat stress tolerance and DNA damage responses **(D).**

Our approach successfully identified genes contributing to stress resilience beyond the well-characterized PQC machinery. Due to functional redundancy within stress response networks, traditional single-gene knockout studies often fail to reveal phenotypic consequences. However, by examining mutant libraries in backgrounds lacking key PQC components, we unearthed previously unrecognized fitness determinants under specific stress conditions. For instance, we identify CanA as a critical factor selective for canavanine stress, likely through acetylation of the unnatural amino acid and a neighboring predicted deacetylase that limits this protective activity. We also reveal the importance of Dps in both heat and oxidative stress in all strains regardless of PQC status, highlighting the ability of this method to uncover stress-specific genes that are otherwise masked in wild-type conditions.

We observed that many genes upregulated in response to stress do not necessarily contribute to stress tolerance under normal conditions. Instead, their importance becomes evident when key stress response genes are absent. This was particularly apparent in the case of oxidative stress, where genes involved in DNA repair became critical only in the absence of the Lon protease. This suggests that transcriptional upregulation does not always equate to functional necessity but may serve as a compensatory mechanism only required under specific genetic perturbations.

Our findings suggest that cellular sensitivity to stress is likely due to the selective loss of specific crucial proteins particular to the state of the cell during that stress rather than a general proteotoxic effect. Heat-induced protein aggregation sequesters key proteins required for survival under stress. For example, we found that the absence of ClpB led to persistent aggregation of RecA, which becomes lethal in a polA- deficient background or when cells immediately encounter genotoxic conditions during heat stress. Extending this result suggests that the detrimental effects of a stress may primarily arise from the loss of specific proteins whose absence becomes particularly deleterious under defined genetic or environmental contexts, rather than a general overall loss in fitness.

In summary, our study underscores the power of using systematic mutagenesis in perturbed backgrounds to elucidate hidden stress response mechanisms. By identifying novel stress determinants, decoupling gene expression from functional necessity, and linking stress sensitivity to the selective loss of key proteins, we provide a refined understanding of bacterial stress responses that can inform future studies in both fundamental and applied microbiology. For example, similar approaches with antibiotic stresses and known resistance genes will allow us to identify additional pathways to target, which is especially important given the alarming rise of antibiotic resistance in many human pathogens.

### Limitations of the study

Here, we show a targeted transposon mutagenesis study in perturbed PQC backgrounds to uncover hidden fitness determinants and novel interactions. However, Tnseq has intrinsic limitations, such as small genes, essential genes with dispensable domains, and nucleoid-associated proteins^44–47^. These issues can result in false positive or negative results. Additionally, it is only possible to study interactions involving genes that can be knocked out.

Another limitation is potential species-specific effects. In this study, we have only examined the phenotype of our model organism, *Caulobacter crescentus*. The heat and oxidative stress sensitivity of PolA has been shown in *E. Coli*^35^, and the heat-stress sensitivity of Δ*clpB* is conserved^16,17,48,49^, but it is possible that the PolA-ClpB synergistic phenotype is Caulobacter-specific.

Finally, we provided mechanistic evidence for PolA’s heat-stress phenotype and its synergistic relationship with ClpB, including RpoH overexpression and RecA aggregation. However, further studies are still needed to explain the deeper mechanistic reasons why RpoH overexpression causes a fitness defect in the Δ*polA* background and the exact function of PolA that leads to these defects.

## Materials availability

Materials generated in this study are listed in the key resources table and available from the lead contract.

## Data and code availability

- RNAseq data have been deposited at GEO at GEO:TBD accession number and are publicly available as of the date of publication.
- RNAseq alignment and annotation pipeline can be found in our GitHub repository. (https://github.com/baldikacti/chienlab-rnaseq)
- Tnseq alignment and annotation pipeline can be found in our GitHub repository. (https://github.com/baldikacti/chienlab-tnseq)
- A web-based application to explore Tnseq data results is hosted from the UMass BMB department’s server. (https://chienbrowser.biochem.umass.edu/chienlab/chienlab_prod2021/)
- Any additional information required to reanalyze the data reported in this paper is available from the lead contact upon request.

## Acknowledgements

This work was supported by National Institutes of Health grants R35GM130320 (PC) and R01GM135931 (PF). B.A was supported in part through the UMass NIH Chemistry and Biology Interface Training Program (T32GM008515). We thank the Jonas lab for sharing strains and plasmids used in this study.

We also thank the Deutchbauer lab for sharing the E. Coli strain (APA766) used in the construction of the Tn-seq library.

## Author contributions

Conceptualization, B.A. and P.C.; methodology, B.A. and P.C.; Investigation, B.A., H.P., V.S., R.Z., P.F., and P.C.; writing—original draft, B.A. and P.C; writing—review & editing, B.A., H.P., V.S., R.Z., P.F., and P.C.; funding acquisition, P.F. and P.C.; resources, P.C.; supervision, P.F., and P.C.

## Metarials and Methods

### Caulobacter crescentus strains

Bacterial strains and plasmids used in this study are listed in the key resources table. Every mutant Caulobacter strain generated from the original NA1000 (WT) strain and grown in PYE medium (2g/L peptone, 1g/L yeast extract, 1 mM MgSO4, and 0.5 mM CaCl2) with aeration at 30°C unless otherwise noted. Antibiotic selections performed at the following concentrations (agar-liquid); kanamycin (25-5 µg/ml), gentamycin (5-0.5 µg/ml), tetracycline (2-1 µg/ml), spectromycin (100-25 µg/ml), streptomycin (5-5 µg/ml), chloremphenicol (1-1 µg/ml).

For plasmid generation, TOP10 *Escherichia coli* competent cells were grown in LB broth (10g/L NaCl, 10g/L tryptone, 5g/L yeast extract) and supplemented with kanamycin (50 µg/ml), gentamycin (40 µg/ml), or spectromycin (50 µg/ml), streptomycin (100 µg/ml) when necessary.

### Cloning and strain construction

Single gene knockouts performed by two-step recombination method with sucrose counter selection using pNTPS138 plasmid^60^. To make *ΔpolA* and *ΔkatG* strains, pNTPS138 plasmid digested using HindIII and EcoRI restriction enzymes and subsequently gel purified. Around 1000 bp upstream and downstream (5’UTR & 3’UTR) regions of the *polA* or *katG* gene and the GENT resistance cassette were PCR amplified. The pNTPS138 plasmid and amplified fragments combined using Gibson Assembly and transformed into TOP10 E.Coli competent cells. The resulting plasmid transformed into electrocompotent NA1000 (*WT*) strain and primary selection perform in PYE agar supplemented with kanamycin. Colonies from the primary selection grown overnight in PYE and plated on PYE agar supplemented with %3 w/v sucrose and gentamicin for secondary selection. Finally, colonies from the secondary selection plated on both PYE agar supplemented with kanamycin or gentamicin, and only colonies which grew on gentamicin but not kanamycin plates were harvested as potential *ΔpolA* or *ΔkatG* strains.

All the double knock-out strains were generated using *θ*cr30 phage transduction as described before with slight modifications^61^. Phage lysates from the donor strains obtained by mixing the 0.5ml of the donor strain at three different concentration of phage at; 50, 5, and 0.5µl. After 15 minutes of incubation at room temperature, cell:phage mixture added to 4ml of molten soft PYE agar (%0.3) and poured on a plate with solidified PYE agar (%1.5). After overnight incubation, soft agar scraped and resuspended with 5 ml of PYE, rigorously vortexed for 2 minutes, 0.1ml of Chloroform added, and centrifuged at 8000g for 30 minutes to get rid of remaining donor cells. The supernatant stored at 4°C as the phage lysate for further use. To move the mutation from donor strain to receiver strain, 1 ml of the phage lysate from the donor strain UV treated at 1.2 joules and mixed with the receiver strain at 1:4 ratio. (50:200µl) The mixture incubated at room temperature for 1 hour and spread on PYE agar plates supplemented with gentamicin, tetracycline, or chloramphenicol.

### Transposon mutagenesis library generation

Transposon mutagenesis libraries used in this study were generated as previously described^20,62^. Briefly, *E.coli* cells containing randomly barcoded EZTn5 plasmids (APA752, gift from Deutschbauer lab) are conjugated with *WT, Δlon, ΔclpA, ΔclpB, and dnaK-NI* (Xylose-inducable *dnaK*) *Caulobacter crescentus* cells separately. *E.coli* donors are kanamycin-resistant and diaminopimelate (DAP) auxotrophs and require it in the media to grow. For conjugation, E*.coli* donor cells Caulobacter strains were mixed at a 1:10 ratio overnight on a PYE agar plate supplemented with DAP (300 uM). The next day, the conjugate was scraped, resuspended, and spread over 14 large (150 x 15 mm) PYE agar plates supplemented with kanamycin (25ug/ml) and without DAP per strain. In this culture, the donor cells will not survive due to no DAP, and acceptor Caulobacter cells will be selected for the Tn5 plasmid due to kanamycin selection.

Colonies were scraped, pooled, and frozen in PYE + 10% glycerol in 1 ml aliquots. For stress condition experiments, 1 aliquot per replicate per strain was thawed in 3.5 ml of PYE or PYE+%0.2 xylose and recovered overnight in a 30*^◦^*C shaker. For all *dnaK-NI* experiments, cells were recovered at saturating xylose concentrations (PYE+%0.2 xylose), and the stress experiments were done at minimal xylose concentrations. (PYE+%0.002) All conditions were performed in quadruplicates, and optical density (OD) measurements were taken at 600nm. Experiments were done in multiple batches.

### Transposon mutagenesis experiments

#### Control environment

Libraries were back diluted to OD 0.008 into 7 ml of PYE or PYE+0.002% xylose and grown overnight until they reached saturation at OD ∼1.6.

#### Heat stress

Libraries diluted to OD of 1 and heat-stressed at low, medium, or high (37, 42, 43.8^◦^C, respectively) for 45 minutes in a Biorad Thermocycler. After 45 minutes, cells diluted back to final OD of 0.008 in 7 ml media for 24-hour growth.

#### Oxidative stress

Libraries were directly diluted back to OD of 0.008 in 7 ml media that contains low, medium, or high (0.025mM, 0.05mM, 0.1mM) level hydrogen peroxide. Cells were grown for 24 hours in these chronic stress conditions.

#### Canavanine stress

Libraries were directly diluted back to OD of 0.008 in 7 ml media that contains low, medium, or high (25ng/ml, 50ng/ml, 100ng/ml) level hydrogen peroxide. Cells were grown for 24 hours in these chronic stress conditions.

### Transposon sequencing PCR library preparation and sequencing

The sequencing libraries for Next-generation sequencing were prepared using a customized three-step PCR protocol (PCR1-2-3). Genomic DNA input was normalized to a concentration of 100 ng/μl. The initial transposon junction amplification (PCR1) was performed using an arbitrary PCR amplification method. This step utilized a forward primer designed (PCR1_F_RBTn5) to align with one end of the transposon and three reverse arbitrary primers (PCR1_R_arb1-2-3). The PCR1 process employed a 2-step cycling protocol with annealing temperatures set at 42°C and 58°C and the number of cycles at 6 and 15, respectively. Subsequently, the second PCR step (PCR2) involved the addition of 16S adapters for Illumina indexing, along with unique molecular identifiers (UMI), and amplification of the library for 36 cycles. Post-PCR2 cleanup was performed using Aline PCRClean DX magnetic beads. The final PCR step (PCR3) incorporated NexteraXT dual indexes in accordance with the manufacturer’s protocol. Post-indexed library (PCR3) cleanup was performed using Aline PCRClean DX magnetic beads.

### Transposon sequencing data analysis

DNA libraries were sequenced in 5 batches with Illumina NextSeq 500 instrument using the NextSeq High output v2.5 single-end 75bp kit. Resulting *fastq* files pruned using a static region from the transposon (TGTATAAGAG) with *seqkit*^54^, PCR duplicates removed with *JE*^53^, reads aligned to NA1000 (NC011916.1) reference genome with *bwa-mem*^63^, and genome positions assigned to the 5’ positions of transposon insertions with *bedtools genomecov*^64^. Subsequently, *bedtools map* used to count either the total or unique transposon insertion counts with either *bedtools map -o sum* or *bedtools map -o count*. For mapping, we used a reference file in which the genes were clipped by 10% from the N and C terminal regions to eliminate spurious transposon insertions in the terminal regions. A snakemake^58^ pipeline of the Tnseq preprocessing/alignment/mapping is available at our GitHub repository. (https://github.com/baldikacti/chienlab-tnseq)

We have applied ComBat-Seq^65^ batchcorrection to the raw counts to minimize the batch effects introduced by the separate sequencing runs.

All the analysis and plotting performed using R Statistical Software (R Core Team 2021).

### RNA sequencing experiments

RNAseq experiments were performed in chronic heat-stress conditions in 96-well plates in a microplatereader. *WT*, *ΔclpB*, and *ΔpolA* strains grown overnight in PYE and back-diluted to 0.1 OD in the morning. Back-diluted cells subsequently plated in a 96-well plate in triplicates (200µl/well) where each replicate was plated to 9 different wells each to get enough final volume. Cells were grown in the plate reader until the mid-log phase at 30°C or 39°C. Finally, the wells with the replicates combined separately, pelleted, and snap frozen.

### RNA sequencing and analysis

Cell pellets from the RNAseq experiments sent to SeqCenter, LLC for RNA isolation. library preparation, and sequencing. Library preparation was performed using Illumina’s Stranded TotalRNA Prep Ligation with Ribo-Zero Plus kit with custom primers for depleting Caulobacter rRNA. Sequencing was done on a NovaSeq X Plus, producing paired end 150bp reads. Demultiplexing, quality control, and adapter trimming was performed with bcl-convert (Illumina).

RNAseq read alignment and mapping was done using a custom pipeline. Briefly, read quality control and filtering performed using *fastq*^56^, reads aligned to reference genome with *bwa-mem*^63^, read quantification performed with *rsubread*^57^, bigwig files generated for manual visualization in genome browsers using *deeptools*^55^, and finally, principal component analysis (PCA) and differential expression analysis are performed using *DESeq2*^66^. The custom pipeline is available at https://github.com/baldikacti/chienlab-rnaseq.

All the analysis and plotting performed using R Statistical Software (R Core Team 2021).

### Growth curve experiments

Growth curve experiments were carried out using a 96-well in a BioTek Epoch microplate reader (Gen6 software). Overnight or log phase cultures are initially diluted to OD 1 for acute stresses and subsequently diluted to OD 0.008 in 96-well plates (in 200 µl) with inducers, antibiotics, and stresses, as needed. The plates were cultured at 30°C and continuous linear shake (330 ppm) for 24 hours unless otherwise noted. Measurements were taken at OD600 every 20 minutes. All experiments performed in biological replicates.

### Spot titration experiments

Overnight or log phase cultures were initially diluted to OD 1 for acute stresses and subsequently diluted to OD 0.1 and made 8 10x titrations in a 96-well plate. 3 µl aliquots from serial dilutions were spotted on PYE agar plates using a multi-channel pipette. PYE agar plates contained inducers, antibiotics, and stresses, as needed. Spot plates were cultured in a 30°C incubator for 2-3 days and imaged using a Syngene G:Box.

### Competition experiments

For competition experiments, the overnight cultures of *WT* and Δ*recA* strains mixed with *WTvenus* strain and Δ*lon* and Δ*lon*Δ*recA* strains mixed with Δ*lonvenus* strain at a 1:1 ratio. The mixtures were inoculated into 5 ml of PYE media at a final OD of 0.0004 to allow for ∼12 doublings. The mixtures cultured in a shaking incubator for 24 hours. The initial population and the final population were imaged using phase contrast and fluorescent microscopy. Final ratios were normalized to initial ratios where the competitive index of 1 equals no fitness change. All experiments performed in biological replicates.

### Protein aggregation assay

In vivo protein aggregation assay was performed as described previously with slight modifications^28^. Experiments were performed as one biological replicate from each strain per experiment for three experiments. Overnights from *WT, ΔclpB, ΔpolA, and ΔclpBΔpolA* strains were back-diluted into three 50 ml PYE per strain for each timepoint (prestress, poststress, and 2h) and grown until the mid-log phase. (OD600 ∼0.8) 40 ml from each was pelleted at 6000g for 10min at RT. Pellets were resuspended in 1 ml PYE, and poststress and 2h conditions were heat-stressed at 44°C for 15 minutes in a tube heater. 2h condition was reinoculated in 50 ml PYE and cultured for 2 hours after stress in a shaking incubator at 30°C. All following steps were performed at 4°C and buffers used during or after cell lysis were supplemented with complete ULTRA protease inhibitor cocktail (Sigma-Aldrich). Samples were put on ice and pelleted. (6000g, 10min) Pellets were resuspended in 1 ml of Buffer A1 (50 mM Tris/HCl pH 8.0, 150 mM NaCl, 12 U/ml Benzonase). Cells were lysed by sonication (50% amp, 10 sec on/off, 6 min on time) Lysates were washed with Buffer A1 2 times after sonication to remove cell debris. (5000g, 10 min) Protein concentration was measured using Bradford assay, and 500 µl of the sample was snap-frozen in liquid nitrogen as the “total protein fraction”. The remaining 500 µl was centrifuged (20000g, 20min) to pellet the insoluble protein fraction. Pellets were resuspended in 300 µl Buffer A2 (50 mM Tris/HCl pH8.0, 150 mM NaCl, %1 Triton X-100), vortexed, sonicated (high, 1 cycle, 30s), and centrifuged (20000g, 20min) for 3 times. Pellets were resuspended in 100 µl of Buffer A2, vortexed, and incubated on ice for 1 hour. After a final sonication (high, 1 cycle, 30s), the samples were snap-frozen as the “insoluble protein fraction”.

### Ponceau and western blotting

24 µl of each protein fraction was mixed with 6 µl of 5x SDS loading dye supplemented with 5% (v/v) β-mercaptoethanol. Protein concentrations were normalized with 1x SDS loading dye relative to the sample with lowest concentration based on the Bradford assay. 10 µl of normalized samples were loaded into 12% SDS-PAGE gel after boiling for 15 min and centrifuging at max for 10-15 min. Separated proteins were transferred to nitrocellulose blotting membranes (Cytiva Amersham Protran 0.2 µm) with a semi-dry transfer unit (Hoefer). The membranes were stained by immersing in Ponceau staining solution and incubating for 20min at RT. The membranes were then washed in ultrapure water before detecting the bands with gel imager (Syngene). Destaining was done by washing the membranes with TBS-T for four times (5min). For western blotting, the membranes were blocked with 5% milk for 1h at RT, washed with TBS-T for three times, and incubated with 1:5000 anti-RecA antibody (Abcam, #ab63797) overnight at 4°C. After washing three times with TBS-T, 1:10000 IRDye 800CW goat-anti-rabbit IgG secondary antibody (LI-COR, #926-32211) was incubated for 1h at RT. Signals were detected by LI-COR Odyssey CLx imager after three TBS-T washes.

