## Supplementary Material for "Stress testing reveals selective vulnerabilities in protein homeostasis"

### Supplementary figures

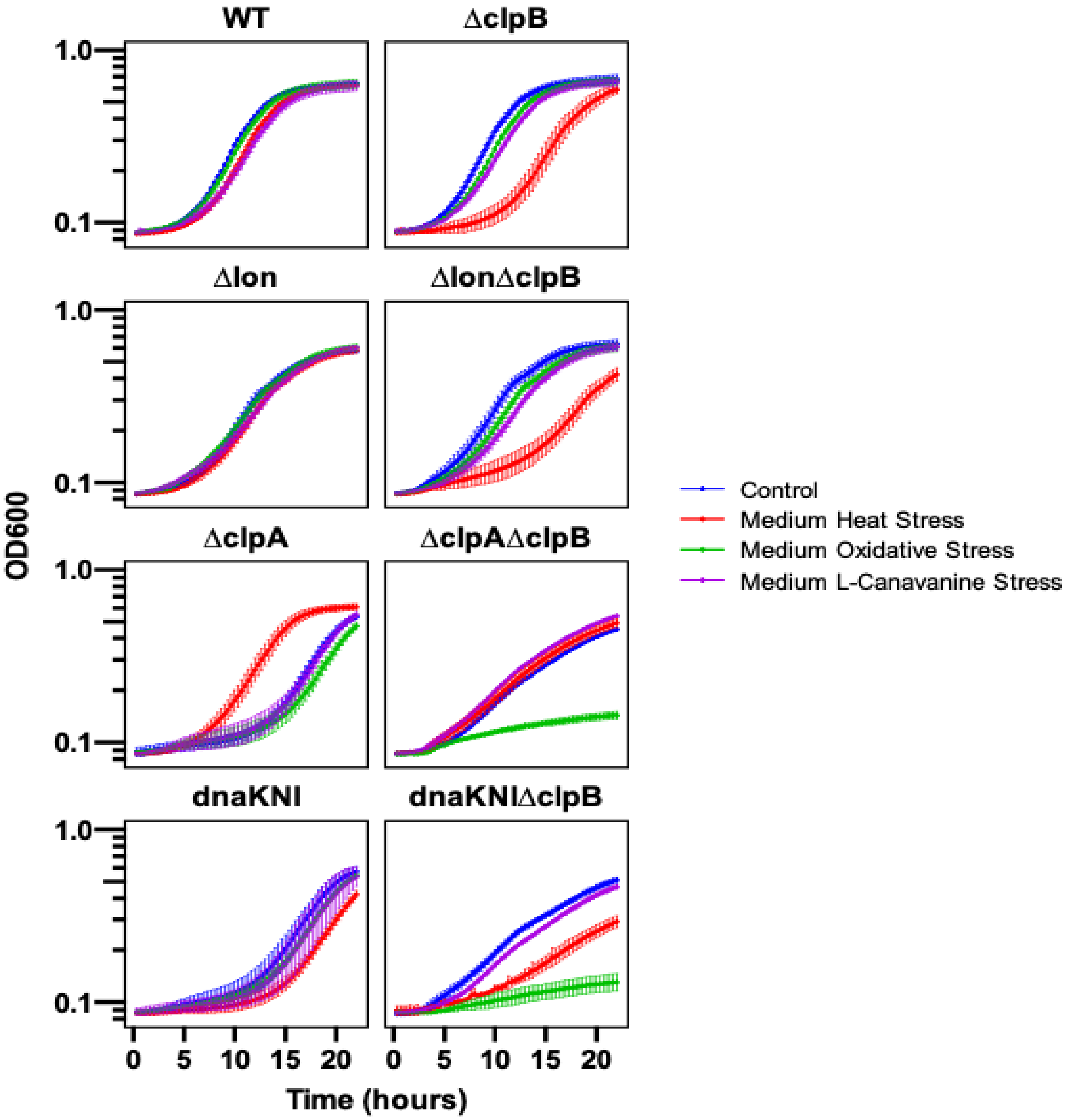


**Supp. Figure 1. Loss of ClpB causes heat-stress sensitivity in *WT, Δlon, ΔclpA*, and *dnaK-NI* backgrounds, related to Figure 1**

Growth curves generated from biological replicates. Conditions tested were; medium heat stress (42°C, 45min), medium oxidative stress (0.05mM $H_2O_2$, chronic), and medium L-Canavanine stress (0.05µg/ml, chronic) . OD600 was measured every 20 minutes in an Epoch2 microplate reader with a continuous linear shake (330ppm). Data is shown as mean +- standard deviation.

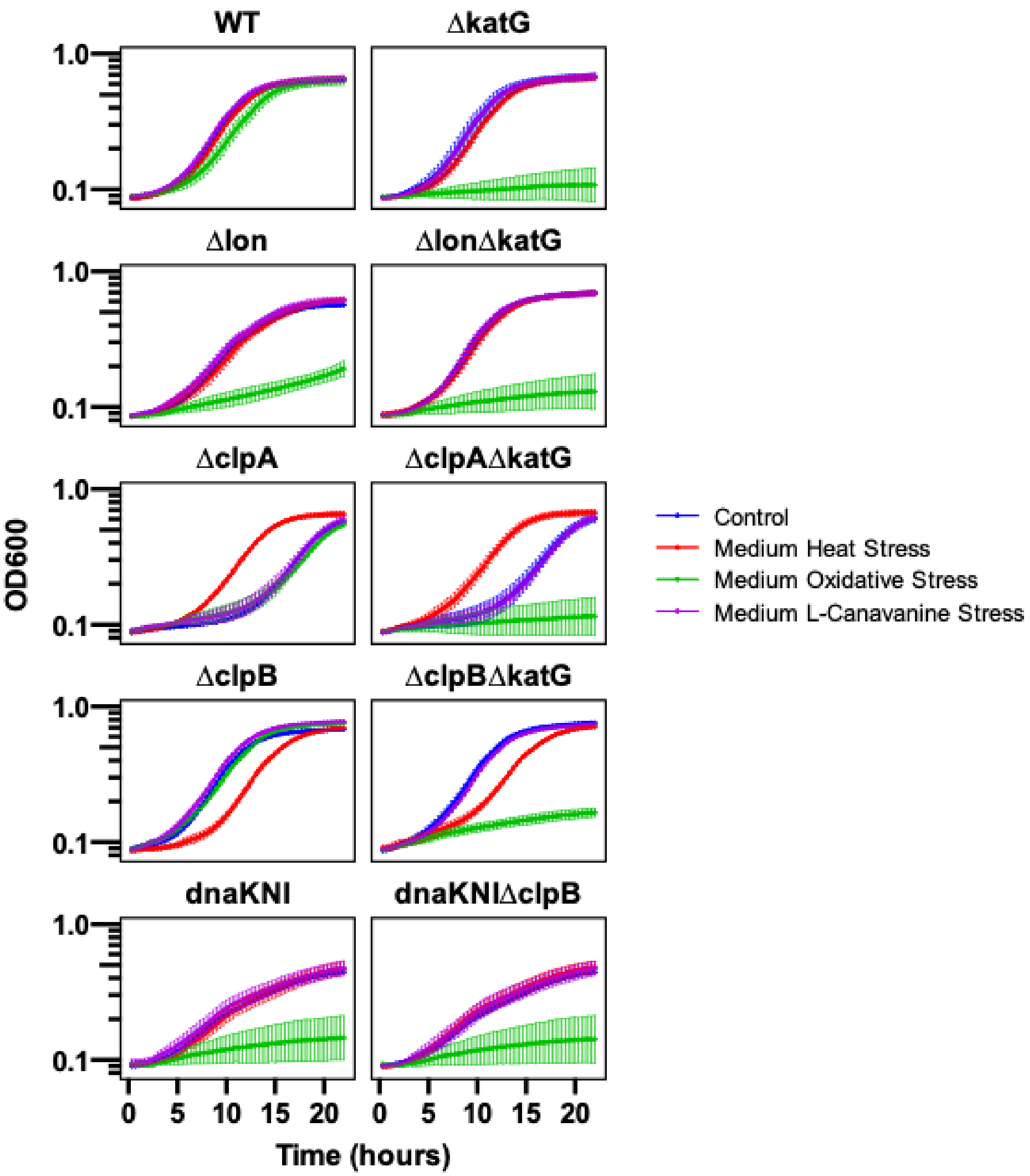


**Supp. Figure 2. Loss of katG causes oxidative-stress sensitivity in *WT, ∆lon, ∆clpA, ∆clpB*, and *dnaK-NI* backgrounds, related to Figure 1**

Growth curves generated from biological replicates. Conditions tested were; medium heat stress (42°C, 45min), medium oxidative stress (0.05mM $H_2O_2$, chronic), and medium L-Canavanine stress (0.05µg/ml, chronic) . OD600 was measured every 20 minutes in an Epoch2 microplate reader with a continuous linear shake (330ppm). Data is shown as mean +- standard deviation.

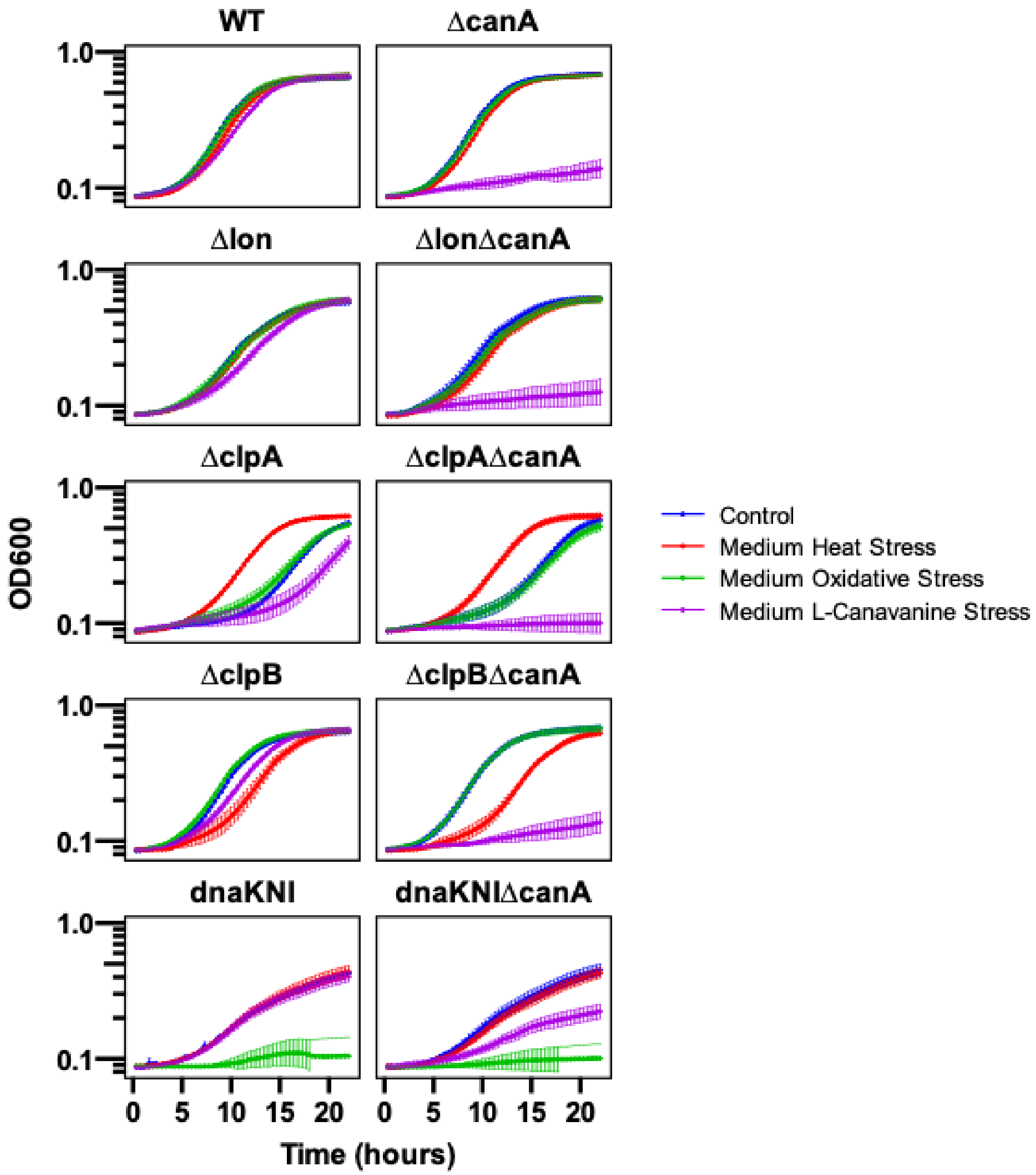


**Supp. Figure 3. Loss of CanA causes L-Canavanine sensitivity in *WT, Δlon, ΔclpA*, and *dnaK-NI* backgrounds, related to Figure 1**

Growth curves generated from biological replicates. Conditions tested were; medium heat stress (42°C, 45min), medium oxidative stress (0.05mM $H_2O_2$, chronic), and medium L-Canavanine stress (0.05µg/ml, chronic) . OD600 was measured every 20 minutes in an Epoch2 microplate reader with a continuous linear shake (330ppm). Data is shown as mean +- standard deviation.

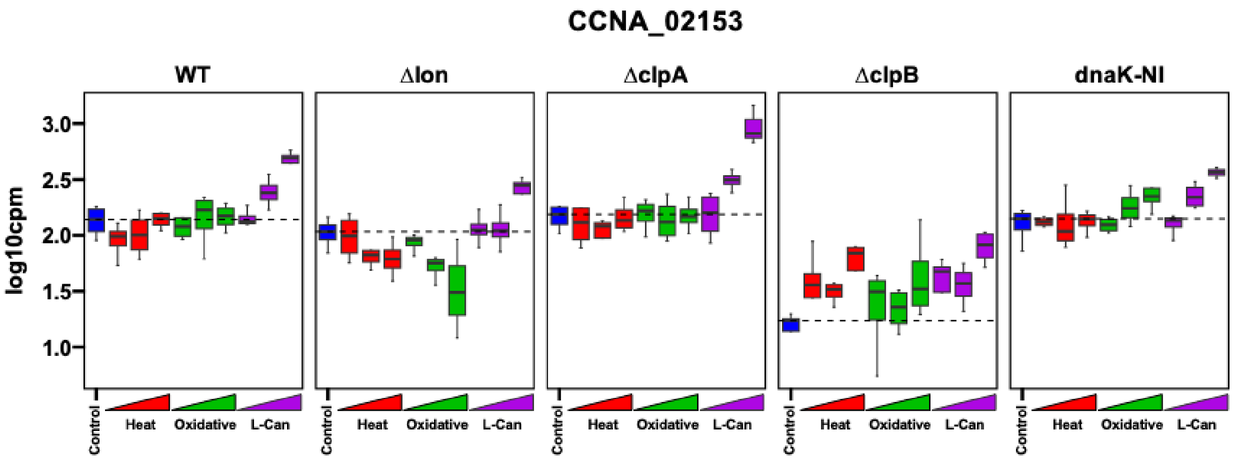


**Supp. Figure 4. Tnseq suggests loss of CCNA_02153 causes protection against L-Canavanine stres in *WT, Δlon, ΔclpA*, and *dnaK-NI* backgrounds, related to Figure 1**

CCNA_02153 transposon insertion profile across all tested transposon mutagenesis libraries and stresses. Insertion counts are presented as log10 batch-corrected counts per million. Stress conditions are color-coded, depicting low, medium, and high stress levels through the triangle. Bars show the median and standard deviation of the insertion counts.

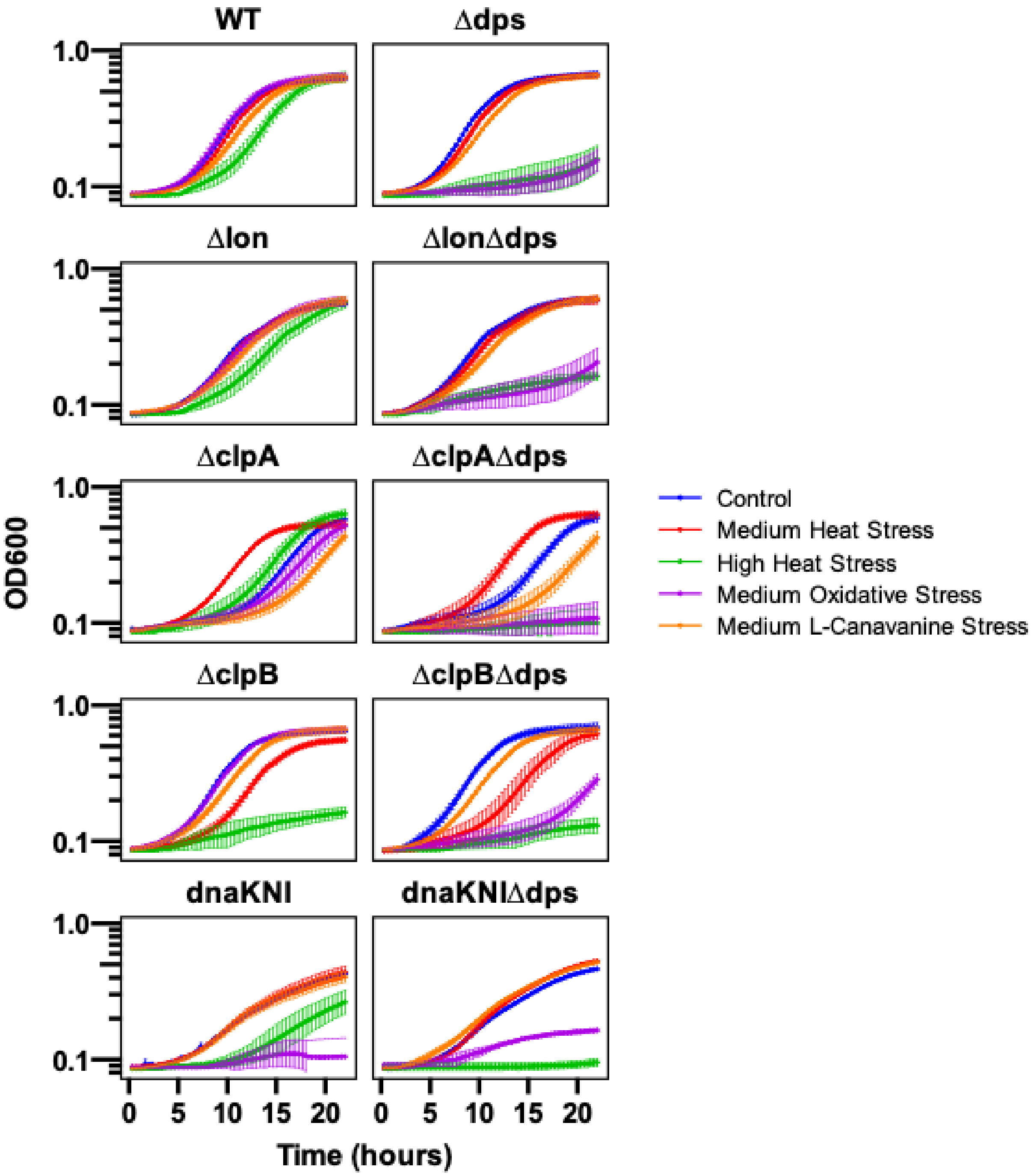


**Supp. Figure 5. Loss of dps causes sensitivity to oxidative and high heat stress in *WT, Δlon, ΔclpA,* and *dnaK-NI* backgrounds, related to Figure 1**

Growth curves generated from biological replicates. Conditions tested were; medium heat stress (42°C, 45min), high heat stress (43.8°C, 45min), medium oxidative stress (0.05mM $H_2O_2$, chronic), and medium L-Canavanine stress (0.05µg/ml, chronic) . OD600 was measured every 20 minutes in an Epoch2 microplate reader with a continuous linear shake (330ppm). Data is shown as mean +- standard deviation.

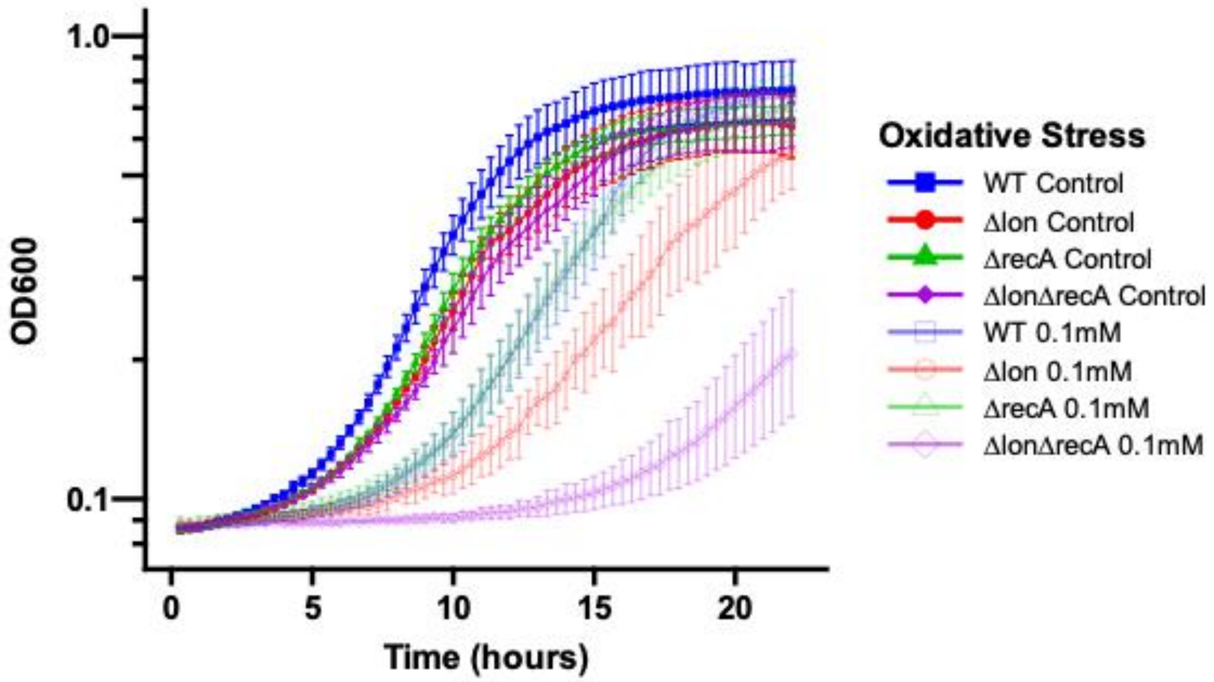


**Supp. Figure 6. Loss of RecA in Δ*lon* background causes a synergistic fitness defect against oxidative stress, related to Figure 2**

Growth curves generated from biological replicates. Conditions tested was chronic 0.1mM $H_2O_2$ stress. OD600 was measured every 20 minutes in an Epoch2 microplate reader with a continuous linear shake (330ppm). Data is shown as mean +- standard deviation.

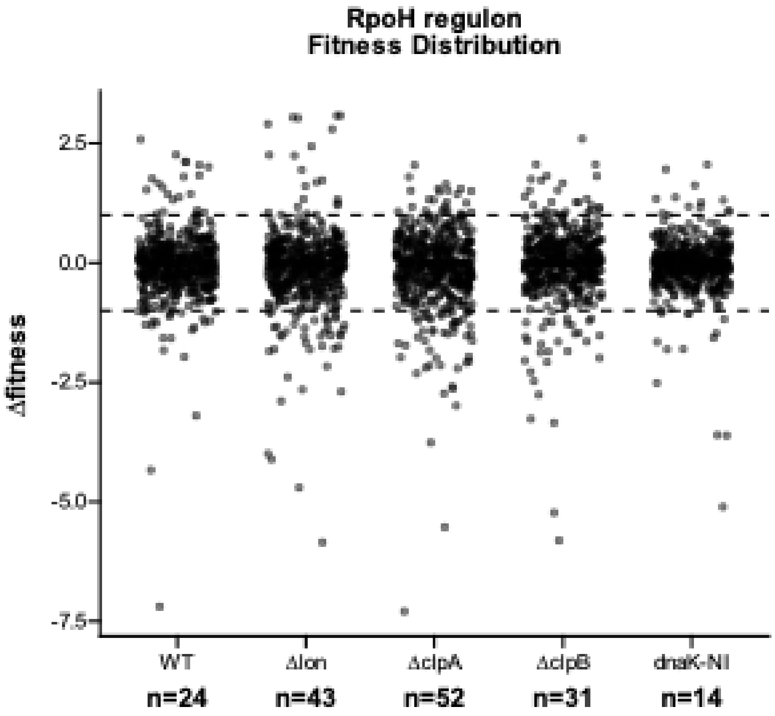


**Supp. Figure 7. Fitness distribution of genes from RpoH regulon, related to Figure 2**

The RpoH regulon obtained from Schramm et al[28]. was used to identify the fitness distribution in our Tnseq data set. Heat stress fitness values are plotted as dots to highlight the distribution between strains. Dashed lines indicate fitness at -1 and +1. The number of genes with less than -1 fitness is labeled on the x-axis.

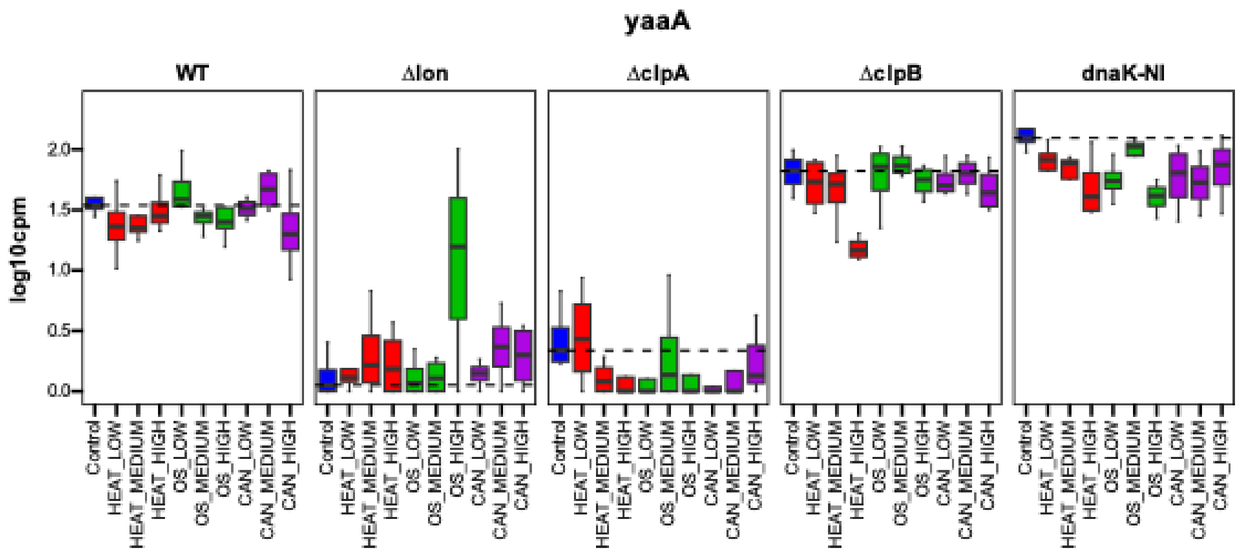


**Supp. Figure 8. Tnseq suggests loss of yaaA causes high heat stress sensitivity in Δ*clpB* background, related to Figure 2**

yaaA transposon insertion profile across all tested transposon mutagenesis libraries and stresses. Insertion counts are presented as log10 batch-corrected counts per million. Stress conditions are color-coded, depicting low, medium, and high stress levels through the triangle. Bars show the median and standard deviation of the insertion counts.

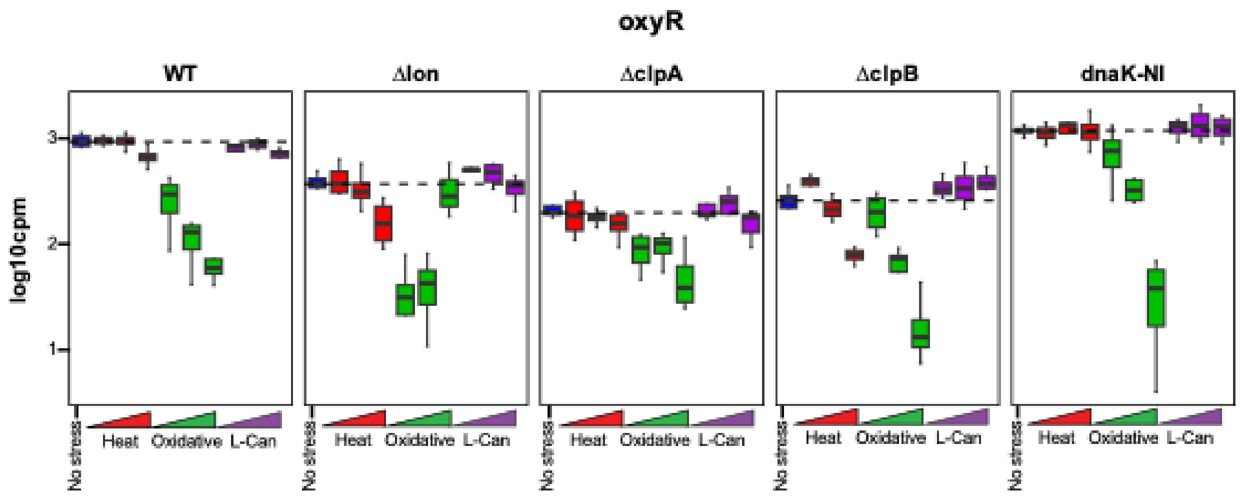


**Supp. Figure 9. Tnseq suggests loss of oxyR causes heat stress sensitivity in Δ*clpB* background, related to Figure 2**

oxyR transposon insertion profile across all tested transposon mutagenesis libraries and stresses. Insertion counts are presented as log10 batch-corrected counts per million. Stress conditions are color-coded, depicting low, medium, and high stress levels through the triangle. Bars show the median and standard deviation of the insertion counts.

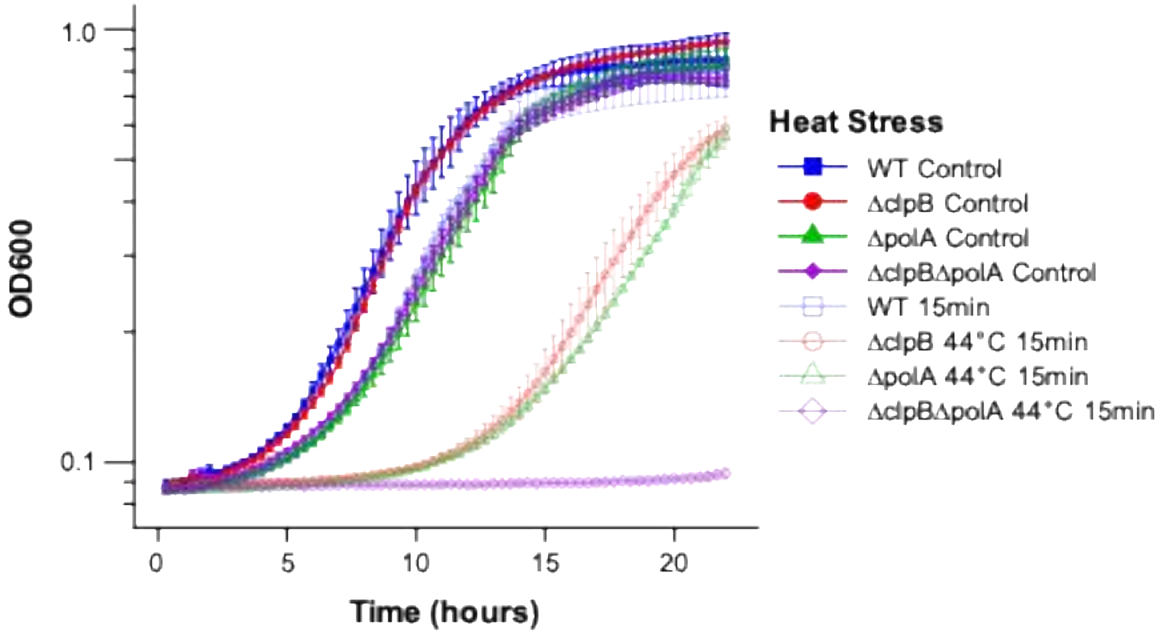


**Supp. Figure 10. Loss of polA causes a synergistic heat stress sensitivity in Δ*clpB* background, related to Figure 3**

Growth curves generated from biological replicates. Conditions tested was heat stress at 44°C for 15 minutes. OD600 was measured every 20 minutes in an Epoch2 microplate reader with a continuous linear shake (330ppm). Data is shown as mean +- standard deviation.

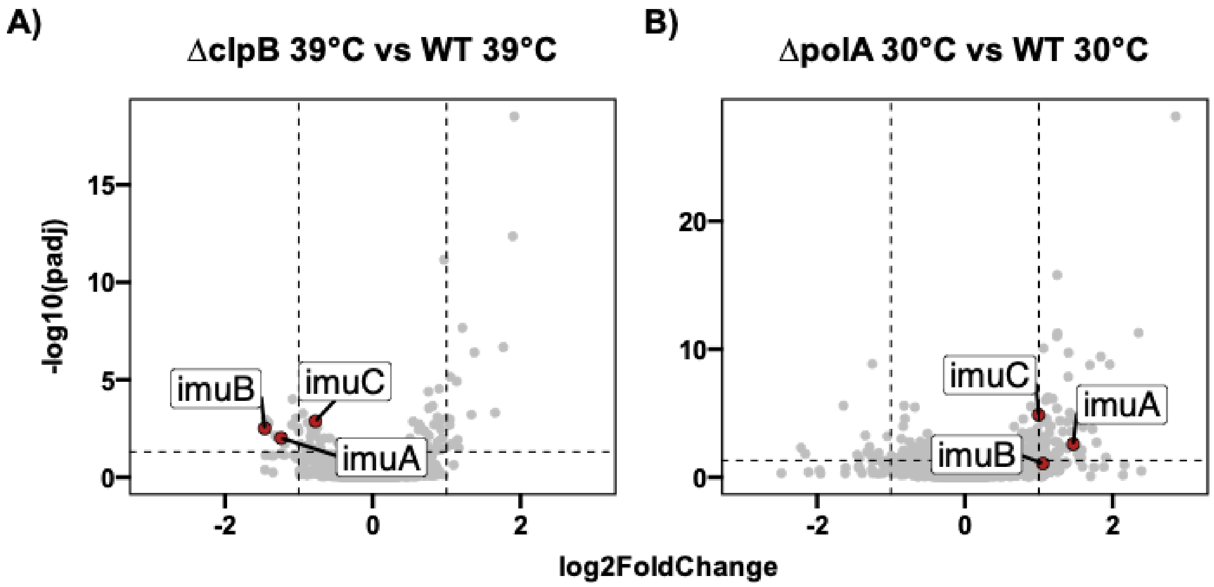


**Supp. Figure 11. Transcriptomic analysis shows the differential regulation of imuABC operon in Δ*clpB* under heat stress and Δ*polA* under regular growth conditions, related to Figure 4**

**(A)** Total RNA sequencing was performed to identify transcriptomic differences between Δ*clpB* (n=3) and *WT* (n=3) strains under heat stress (39°C, chronic) and data collected during mid-log phase growth (~8 hour). Volcano plots show log2 fold change on the x-axis, and -log10 false discovery rate (FDR) on the y-axis. *imuABC* operon genes highlighted on the plot with red dots with black circles. RNAseq experiment performed with biological triplicates. **(B)** Total RNA sequencing was performed to identify transcriptomic profile of Δ*polA* (n=2) strain compared to *WT* (n=3) strain under regular growth conditions, and data collected during mid-log phase growth (~8 hour). Volcano plots show log2 fold change on the x-axis, and -log10 false discovery rate (FDR) on the y-axis. *imuABC* operon genes highlighted on the plot with red dots with black circles. RNAseq experiment performed with biological duplicates or triplicates, as indicated above.

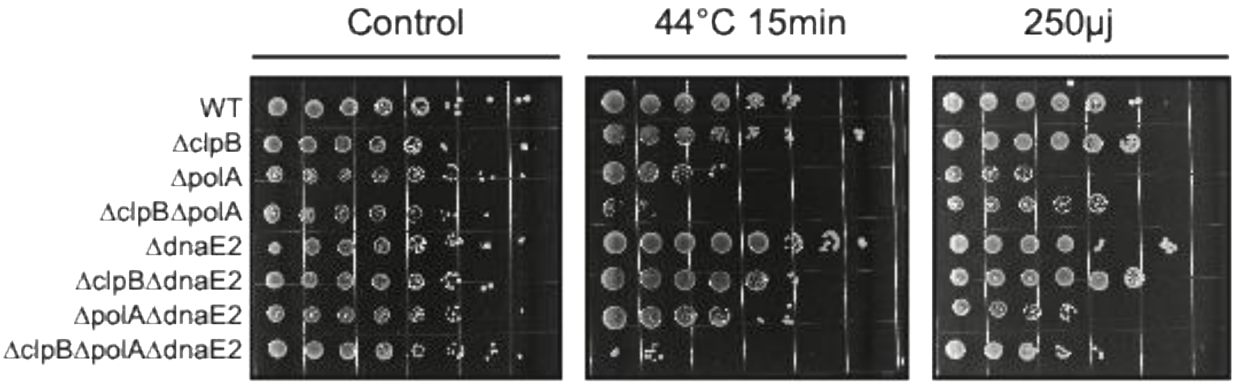


**Supp. Figure 12. Loss of dnaE2 (imuC) does not cause a synergistic fitness effect under heat or oxidative stress with PolA or ClpB, related to Figure 4**

Spot titration (10-fold dilution series) of *WT, ΔclpB, ΔpolA, ΔclpBΔpolA, ΔdnaE2, ΔclpBΔdnaE2, ΔpolAΔdnaE2*, and *ΔclpBΔpolAΔdnaE2* strains under 44°C 15min heat stress, and acute UV stress (250µjoules). Stress conditions applied to cells before spotting, cells washed with PYE once by centrifugation, and spotted on to PYE agar (%1.5) plates.

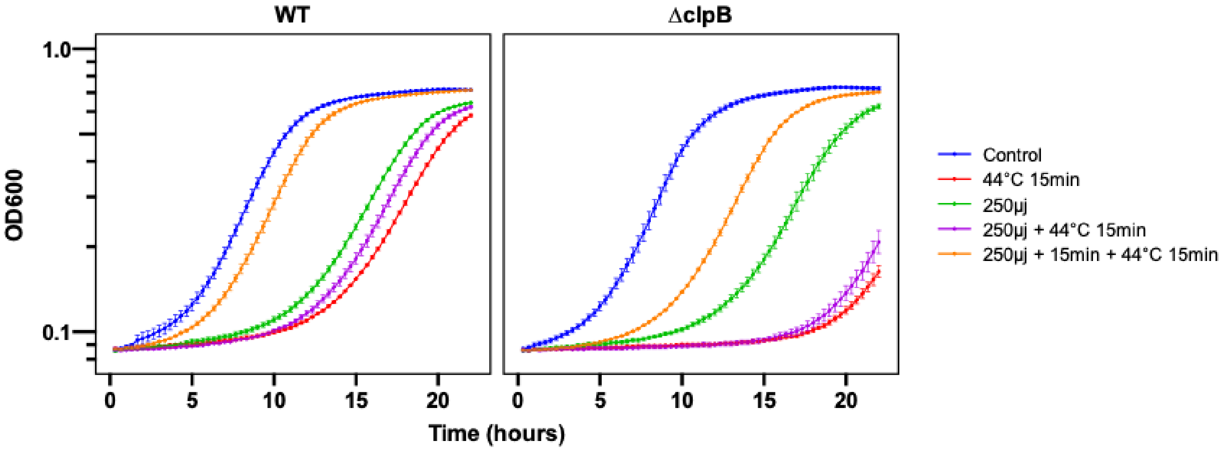


**Supp. Figure 13. Upregulation of SOS response provides protection against heat stress in *WT* and *ΔclpB* strains, related to Figure 4**

Growth curves generated from biological replicates. Conditions tested were;heat stress (44°C 15min), UV stress (250µjoules), heat + UV, and heat + UV stress with a 15 minute recovery period in between stresses. OD600 was measured every 20 minutes in an Epoch2 microplate reader with a continuous linear shake (330ppm). Data is shown as mean +- standard deviation.

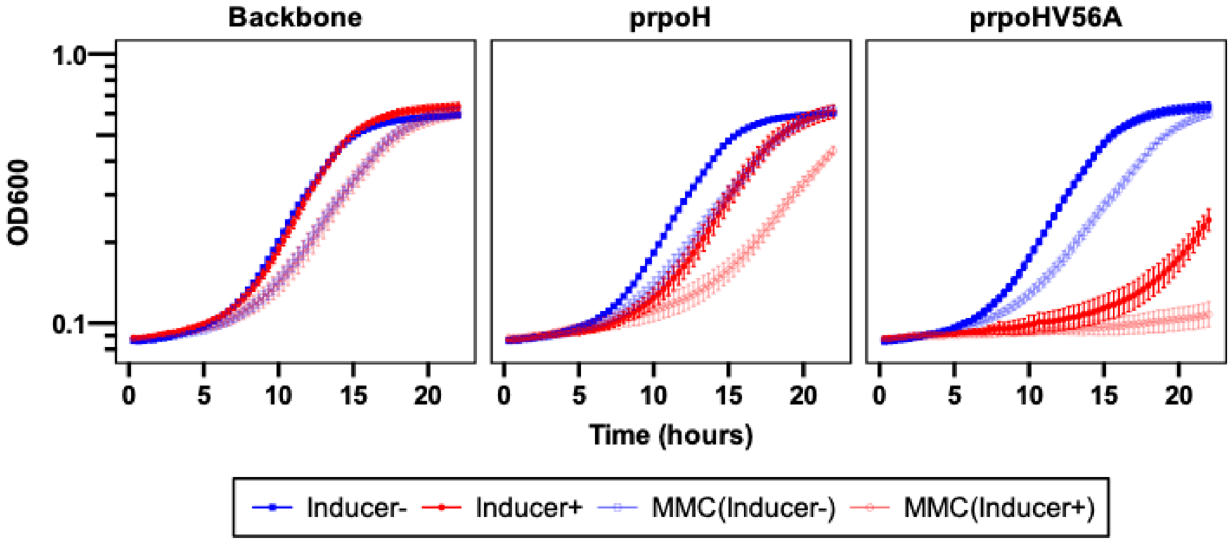


**Supp. Figure 14. RpoH overexpression does not sensitize cells to DNA damage, related to Figure 4**

Growth curves generated from biological replicates. Conditions tested were; MMC stress (0.05µg/ml, chronic) with or without inducer (0.5mM Vanillic acid). Tested strains contained either the plasmid (pJ14) backbone, van-inducible rpoH plasmid, or van-inducible rpoHV56A plasmid. OD600 was measured every 20 minutes in an Epoch2 microplate reader with a continuous linear shake (330ppm). Data is shown as mean +- standard deviation.

#### Supplementary tables

##### Supp. Table 1. Genes that show large fitness change under single stress in all strains

| LOCUS TAG | GENE ID | STRESS |
|---|---|---|
| **ABS(FOLDCHANGE) > 1.5** | | |
| CCNA_01217 | N/A | L-CANAVANINE |
| CCNA_01242 | N/A | L-CANAVANINE |
| CCNA_02154 | canA | L-CANAVANINE |
| CCNA_01070 | N/A | HEAT |
| CCNA_02525 | N/A | HEAT |
| CCNA_03577 | polA | HEAT |
| CCNA_00562 | N/A | OXIDATIVE |
| CCNA_02925 | N/A | OXIDATIVE |
| CCNA_03359 | N/A | OXIDATIVE |
| CCNA_03577 | N/A | OXIDATIVE |
| CCNA_03811 | oxyR | OXIDATIVE |

##### Supp. Table 2. Genes that show large fitness change under two stresses in all strains

| LOCUS TAG | GENE ID | STRESS |
|---|---|---|
| **ABS(FOLDCHANGE) > 1.5** | | |
| CCNA_02966 | dps | HEAT,OXIDATIVE |
| CCNA_03577 | N/A | HEAT,OXIDATIVE |
| **ABS(FOLDCHANGE) > 1** | | |
| CCNA_02966 | dps | HEAT,OXIDATIVE |
| CCNA_03577 | polA | HEAT,OXIDATIVE |
| CCNA_00392 | N/A | HEAT,OXIDATIVE |
| CCNA_02088 | N/A | HEAT,OXIDATIVE |
| CCNA_01217 | N/A | OXIDATIVE,L-CANAVANINE |
| CCNA_01242 | N/A | OXIDATIVE,L-CANAVANINE |
| CCNA_01947 | N/A | OXIDATIVE,L-CANAVANINE |
| CCNA_02941 | greA | OXIDATIVE,L-CANAVANINE |

##### Supp. Table 3. Genes that show large fitness change under two or more strains for all stresses

| LOCUS TAG | GENE ID | STRAINS |
|---|---|---|
| **ABS(FOLDCHANGE) > 1.5** | | |
| CCNA_00001 | N/A | *WT,Δlon* |
| CCNA_00043 | nusA | *WT,ΔclpB* |
| CCNA_00050 | N/A | *WT,dnaK-NI* |
| CCNA_01377 | N/A | *WT,ΔclpB* |
| CCNA_02345 | N/A | *WT,ΔclpB* |
| CCNA_02572 | N/A | *WT,ΔclpA* |
| CCNA_03026 | N/A | *WT,dnaK-NI,Δlon* |
| CCNA_03052 | N/A | *WT,dnaK-NI* |
| CCNA_03729 | N/A | *WT,dnaK-NI* |
| CCNA_R0125 | N/A | *WT,Δlon* |
| CCNA_R0141 | N/A | *WT,ΔclpB,Δlon* |
| CCNA_01254 | smpB | *dnaK-NI,Δlon* |
| CCNA_03256 | N/A | *dnaK-NI,Δlon* |
| CCNA_00285 | N/A | *ΔclpA,Δlon* |
| CCNA_02925 | N/A | *ΔclpB,Δlon* |
| CCNA_03021 | N/A | *ΔclpB,Δlon* |
| CCNA_03420 | N/A | *ΔclpB,Δlon* |
| CCNA_03970 | N/A | *ΔclpB,Δlon* |

**Supp. Table 4. Primers used for transposon mutagenesis library preparation**

| PRIMER NAME | SEQUENCE |
| --- | --- |
| **PCR1** | |
| PCR1_F_RBTn5 | CGCCCTGCAGGGATGTCCACGAG |
| PCR1_R_arb1 | GTCTCGTGGGCTCGGAGATGTGTATAAGAGACAGNNNNNNNNNNACGCC |
| PCR1_R_arb2 | GTCTCGTGGGCTCGGAGATGTGTATAAGAGACAGNNNNNNNNNNCCTGG |
| PCR1_R_arb3 | GTCTCGTGGGCTCGGAGATGTGTATAAGAGACAGNNNNNNNNNNCCTCG |
| **PCR2** | |
| PCR2_F6_Rbjunc2 | TCGTCGGCAGCGTCAGATGTGTATAAGAGACAGNNNNNNGCCGGCCGGT TGAGATGTGTA |
| PCR2_R_universal | GTCTCGTGGGCTCGGAGATG |
| **PCR3** | |
| N701 | CAAGCAGAAGACGGCATACGAGATTCGCCTTAGTCTCGTGGGCTCGG |
| N702 | CAAGCAGAAGACGGCATACGAGATCTAGTACGGTCTCGTGGGCTCGG |
| N703 | CAAGCAGAAGACGGCATACGAGATTTCTGCCTGTCTCGTGGGCTCGG |
| N704 | CAAGCAGAAGACGGCATACGAGATGCTCAGGAGTCTCGTGGGCTCGG |
| N705 | CAAGCAGAAGACGGCATACGAGATAGGAGTCCGTCTCGTGGGCTCGG |
| N706 | CAAGCAGAAGACGGCATACGAGATCATGCCTAGTCTCGTGGGCTCGG |
| N707 | CAAGCAGAAGACGGCATACGAGATGTAGAGAGGTCTCGTGGGCTCGG |
| N708 | CAAGCAGAAGACGGCATACGAGATCCTCTCTGGTCTCGTGGGCTCGG |
| N709 | CAAGCAGAAGACGGCATACGAGATAGCGTAGCGTCTCGTGGGCTCGG |
| N710 | CAAGCAGAAGACGGCATACGAGATCAGCCTCGGTCTCGTGGGCTCGG |
| S501 | AATGATACGGCGACCACCGAGATCTACACTAGATCGCTCGTCGGCAGCG TC |
| S502 | AATGATACGGCGACCACCGAGATCTACACCTCTCTATTCGTCGGCAGCGT C |
| S503 | AATGATACGGCGACCACCGAGATCTACACTATCCTCTTCGTCGGCAGCGT C |
| S504 | AATGATACGGCGACCACCGAGATCTACACAGAGTAGATCGTCGGCAGCG TC |
| S505 | AATGATACGGCGACCACCGAGATCTACACGTAAGGAGTCGTCGGCAGCG TC |
| S506 | AATGATACGGCGACCACCGAGATCTACACACTGCATATCGTCGGCAGCG TC |
| S507 | AATGATACGGCGACCACCGAGATCTACACAAGGAGTATCGTCGGCAGCG TC |
| S508 | AATGATACGGCGACCACCGAGATCTACACCTAAGCCTTCGTCGGCAGCG TC |
| S510 | AATGATACGGCGACCACCGAGATCTACACCGTCTAATTCGTCGGCAGCG TC |
| S511 | AATGATACGGCGACCACCGAGATCTACACTCTCTCCGTCGTCGGCAGCG TC |

**Supp. Table 5. Primers used for strain generation**

| PRIMER NAME | SEQUENCE |
| --- | --- |
| **∆POLA PRIMERS** | |
| 5F_Pol1_pNTPS138 | attgaagccggctggcgccaagcttcggcgtggatcacatagcg |
| 5R_Pol1_pNTPS138 | agcgtcagaccccgtagaaaagatcagaaaccgccgtgcctcagttcctg |
| 3F_pol1_pNTPS138 | cgacccaagtaccgccacctaatccttcgccggtgatcggctcg |
| 3R_pol1_pNTPS138 | gtcacggccgaagctagcgaattcgccaccaagatgaaggtggtcaacaagg |
| **∆KATG PRIMERS** | |
| 5F_katG_pNTPS138 | GAAGCCGGCTGGCGCCAAGCTTGGCTGGCGGCCAGCTGGGAC |
| 5R_katG_pNTPS138 | CAGACCCCGTAGAAAAGATCCTACGACAACCCGACGCGCCTAGAAGCCG |
| 3F_katG_pNTPS138 | gacccaagtaccgccaccTAAgatctggcggcttaggatct |

| | |
|---|---|
| 3R_katG_pNTPS138 | CGTCACGGCCGAAGCTAGCGAATTccaacagccagatcgcccac |
| **∆DPS PRIMERS** | |
| 5F_dps_pNTPS138 | AATTGAAGCCGGCTGGCGCCAAGCTTggaaagatgcgggttcgccagtccagg |
| 5R_dps_pNTPS138 | agcgtcagaccccgtagaaaagatcctttgccgggagcggggatggagaag |
| 3F_dps_pNTPS138 | atcgacccaagtaccgccacctaaccatcccgcgtgcgggcgggggctcgg |
| 3R_dps_pNTPS138 | GCGTCACGGCCGAAGCTAGCGAATTtcgacacccccagcatctgggccgc |
| **∆CANA PRIMERS** | |
| 5F_canA_pNTPS138 | CAATTGAAGCCGGCTGGCGCCAAGCTTcggcgatggtggcgtcagccatgttg |
| 5R_canA_pNTPS138 | agcgtcagaccccgtagaaaagatcggtctaggagtctccgatggctctcgc |
| 3F_canA_pNTPS138 | tatcgacccaagtaccgccacctaagtccctgcgtccttcgaggctcgcc |
| 3R_canA_pNTPS138 | CGTCACGGCCGAAGCTAGCGAATTagacggtctcgtagaggcagttgtgg |
| **GENTR PRIMERS** | |
| F_gentR | gatcttttctacggggtctgacgc |
| R_gentR | ttaggtggcggtacttgggtcgatatc |
